# Conserved transcriptomic heat stress response signatures in coral recruits selectively bred from thermally distinct broodstock in a low-differentiation system

**DOI:** 10.64898/2026.06.23.733312

**Authors:** Richard C. Edmunds, Alex Macadam, Carys A. Morgans, Guy A. McCutchan, Thomas Danhorn, Patrick W. Laffy, Patrick Buerger, Madeleine J.H. van Oppen, Kate M. Quigley, Annika M. Lamb

## Abstract

Thermal history provenancing can guide the choice of parental broodstock for selective breeding of corals from distinct reefs and has been proposed as an intervention for enhancing climate resilience. However, the genetic and molecular mechanisms underlying resultant offspring responses to heat stress, particularly during early life stages, remain poorly understood. Here, we generated *Acropora tersa* larvae and recruits by crossing parental colonies from the historically warmer Martin Reef and cooler Davies Reef and assessed the effects of within- and between-reef crosses on genetic diversity and transcriptional responses to heat stress. Genome-wide single nucleotide polymorphism analyses showed that broodstock from Martin and Davies Reefs were weakly differentiated (*F_ST_* = 0.008) and exhibited comparable heterozygosity, as did all larval offspring groups. Transcriptomic analyses of recruits exposed to heat stress (32 °C for 36 days) revealed that both within- and between-reef offspring groups activated conserved stress-response pathways, with seven genotype-independent heat-responsive genes detected across all offspring groups. Differential expression and enrichment analyses showed induction of defence, protein homeostasis, intracellular transport, and metabolic processes alongside repression of growth- and signalling-related functions, consistent with the Type A General Coral Stress Response. Taken together, these findings suggest that the benefits of thermal history provenancing-informed selective breeding may be limited in low-differentiation systems and that targeted pre-screening of broodstock may help capture functional genetic variation relevant to restoration applications.

## 1. Introduction

The ongoing degradation of coral reefs threatens the livelihoods, food sources, cultures, and economies of human communities and may lead to severe losses of marine biodiversity (IPCC, 2022). The primary drivers of this decline are marine heatwaves, which cause bleaching (i.e., the loss of photosymbionts from coral holobionts) and often kill reef-building corals (Hoegh-Guldberg, 1999). There have now been four global mass coral bleaching events, culminating in the fourth event that began in 2023 and has been driven by increasingly frequent, prolonged, and intense marine heatwaves (Spady et al., 2026). During this period, coral reefs worldwide experienced unprecedented thermal stress (e.g., Neely, et al., 2024), with 46% of reefs on the Great Barrier Reef (GBR) exposed to the highest levels of heat stress on record (Cantin et al., 2024). Despite the severity of these impacts overall, variation in heat tolerance exist within and among coral populations (Denis et al., 2024; Naugle et al., 2024), with some of this variation having a host genetic basis (Dixon, et al., 2015; Howells, et al., 2016). Given time, evolutionary forces such as gene flow and natural selection can act on this variation to increase coral thermal resilience (Torda & Quigley, 2022). However, the current and predicted rates of coral demographic decline on reefs suggest their natural rate of evolution is not keeping pace with climate change (Lachs et al., 2024).

Corals can adapt to their thermal environment, with populations inhabiting reefs with histories of high temperatures often exhibiting comparatively greater thermotolerance (Howells et al., 2016). Warm-adapted corals can be bred with corals from reefs with cooler thermal histories to generate offspring that may have improved heat tolerance relative to corals native to the cooler reefs (Eirin-Lopez & Putnam, 2019; Quigley, 2024). Such thermally enhanced offspring could be deployed onto cooler reefs to enhance resilience of resident populations against future summer heatwaves (van Oppen et al., 2015). This approach applies the concept of thermal history provenancing where future climate-suitable stock is sourced and used to seed sites in anticipation of future climate change conditions (Aitken & Whitlock, 2013; Prober et al., 2015).

Reef thermal histories have guided the selection of source reefs for research programs aiming to enhance offspring heat tolerance (Quigley, et al., 2020a). Results from thermal history provenancing in corals have been mixed, with some but not all offspring families exhibiting enhanced thermal tolerance (Howells et al., 2021; Quigley et al., 2021; Quigley, Randall, et al., 2020b; Macadam et al., 2025). These variable findings are frequently linked to variable levels of genetic differentiation among coral populations (Frankham et al., 2019) as well as thermal tolerance variation within populations (Humanes, et al., 2022; Denis, et al., 2024) and broodstock selection approaches that fail to capture heritable variation in thermal tolerance (Lachs, et al., 2026; Lamb, et al., *in press*). At one end of the spectrum, crossing corals from reefs connected by high gene flow, as is the case amongst GBR reefs for some species (Fuller et al., 2020), is likely to produce offspring that are similar to those from within-reef crosses, limiting the potential for enhancement. At the other end of the spectrum, crosses between corals from reefs with limited gene flow and strong local adaptation may yield offspring with enhanced target traits but also carry a greater risk of disrupting local adaptations and causing outbreeding depression (Frankham et al., 2011).

Genomic technologies such as single nucleotide polymorphism (SNP) sequencing enable the quantification of genetic diversity and differentiation, which provide critical insights into population structure and evolutionary potential that inform assessments of adaptive capacity and selective breeding strategies (Quigley, et al., 2020b; Baums et al., 2022; Torda and Quigley, 2022). Moreover, transcriptomic approaches such as ribonucleic acid sequencing (RNAseq) can be used to understand how organisms respond to environmental pressures and identify the genes and molecular pathways underpinning adaptive resilience (i.e., capacity to persist and adapt under environmental change) and survival (Louis, et al., 2017; Drury, 2020). RNAseq therefore provides a functional complement to SNP analyses without implying direct correspondence between genomic and transcriptomic datasets (e.g., Dziedzic, et al., 2019).

We used the broodstock and within- and between-reef offspring groups of *Acropora tersa* (Rasmussen et al., 2025; previously referred to as *A. hyacinthus* “neat”; Naugle et al., 2024) as described by Macadam et al. (2025). These corals originated from reefs in the Cairns/Cooktown (i.e., northern) and Townsville/Whitsunday (i.e., central) Management Areas of the GBR, which experience contrasting thermal regimes given the warmer and cooler monthly maximum mean (MMM) temperatures, respectively (Smith & Spillman, 2019). Macadam et al. (2025) found that, overall, thermal history provenancing-informed selective breeding between these regions did not consistently influence offspring performance (e.g., growth, survival, or bleaching response) under heat stress conditions. These findings raise the question of whether underlying genetic composition and transcriptional responses differ among within- and between-reef offspring groups in the absence of consistently divergent physiological phenotypes. To address this, we applied genome-wide SNP analyses to quantify genetic diversity and differentiation among broodstock and larval offspring groups and, in parallel, RNAseq to characterise group-level transcriptomic responses of recruits exposed to ambient and heat stress conditions.

## 2. Methods

### 2.1 Broodstock colonies, experimental design, and DNA/RNA sampling

Broodstock colonies of *A. tersa* (Figure 1A) were collected from Martin (−14.7713, 145.3694) and Davies (−18.8385, 147.6324) reefs prior to the December 2021 spawning event (Great Barrier Reef Marine Park Authority permit G20/44541.1; Macadam et al., 2025; Figure 1B). The MMM temperature of a location is commonly used as the baseline against which coral heat stress is measured (Ainsworth et al., 2016); the MMMs of Martin and Davies reefs are 28.63°C and 28.39°C, respectively (www.reefwatch.noaa.gov; Strong et al., 1997). Despite a MMM difference of only ∼0.24 °C between reefs, adult broodstock from Martin exhibited markedly greater thermotolerance than those from Davies, with survival under heat stress (32 °C for 7 days) declining by ∼3% and ∼36% relative to controls, respectively (Macadam et al., 2025). Degree Heating Weeks (DHWs) were calculated following NOAA Coral Reef Watch conventions as the time-integrated accumulation of daily thermal stress exceeding the local bleaching threshold (MMM + 1 °C), expressed in °C-weeks. For each day of the 36-day heat-stress exposure, only temperatures above MMM + 1 °C contributed to DHW accumulation, with the daily excess temperature summed and converted to °C-weeks to represent cumulative thermal stress over the experimental period.

**Figure 1.**
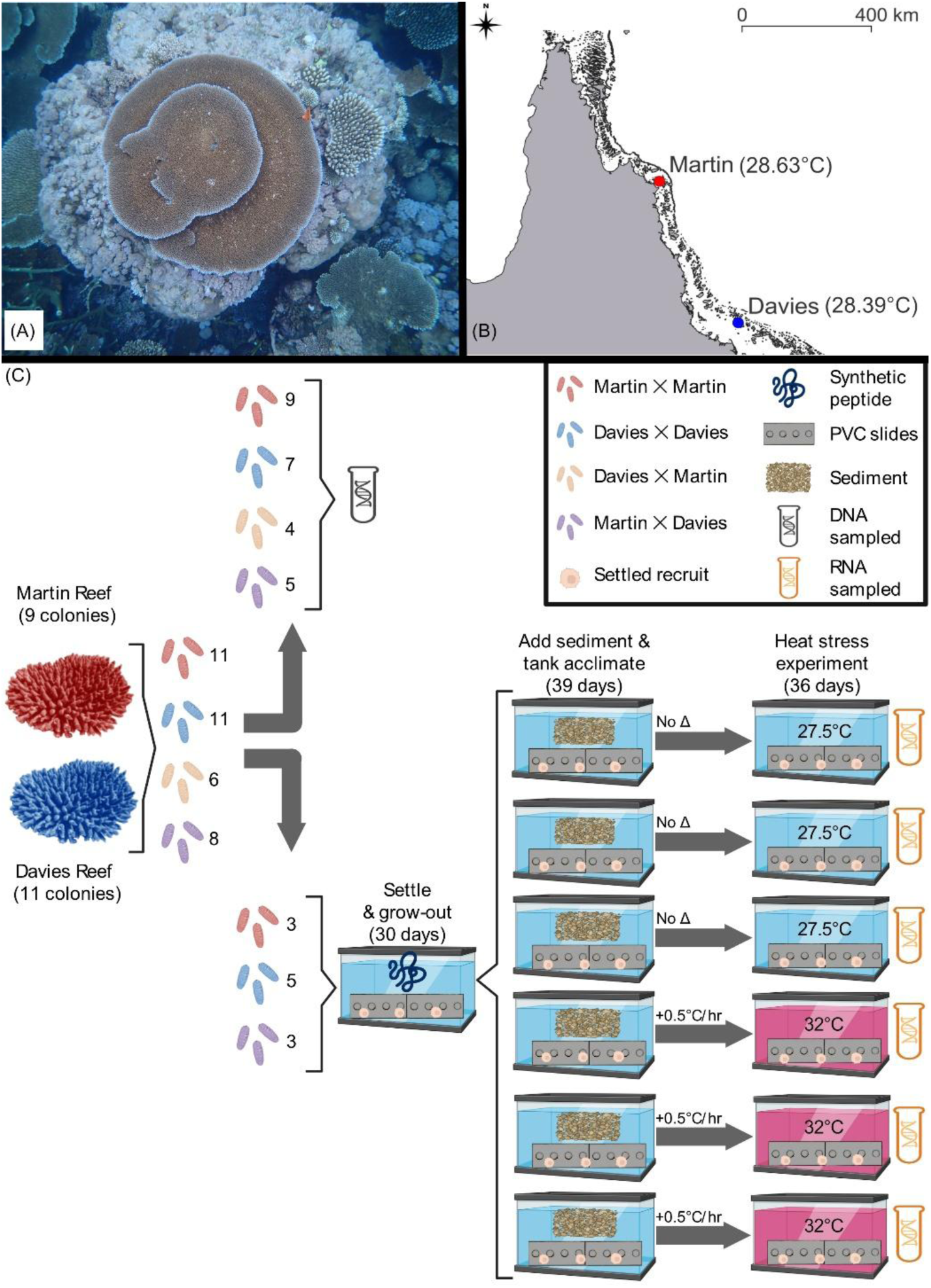
*Acropora tersa* morphology, sampling locations, and experimental design. (A) Photo showing target species *A. tersa*. Photo credit: Véronique Mocellin (AIMS). (B) Map of Queensland Australia showing collection locations for Martin (red) and Davies (blue) broodstock from northern and central regions of the Great Barrier Reef, respectively, with monthly maximum mean temperature for each reef provided in parentheses. (C) Experimental design for larval breeding and recruit heat stress experiment (see also Figure 1 in Macadam et al., 2025). DNA and RNA sampling points are shown as black and orange test tube icons, respectively. Settlement-inducing synthetic peptide (1 µM) was applied during 30-day settlement and grow-out period before 39-day acclimation and Wood Reef sediment exposure. Recruit RNA samples were collected following a 36-day exposure to ambient (27.5°C) and heat stress (32°C) treatment conditions (blue and red tanks), respectively. Numbers refer to the number of families used in each treatment unless otherwise specified. The experimental design diagram (Figure 1C) was created using BioRender (http://BioRender.com).

All broodstock colonies were transferred to the National Sea Simulator (SeaSim) at the Australian Institute of Marine Science (AIMS) and housed in outdoor aquarium systems exposed to natural daylight and moonlight. Coral showing signs of “setting” (Harrison, 2011) were isolated and monitored for spawning. Egg-sperm bundles were collected from spawned colonies, and the eggs and sperm were separated by agitating the bundles on a 120 μm filter in 0.2 μm filtered seawater (FSW). Washed eggs and sperm (sperm density of 1 ✕ 10^6^ cells per mL) from a single dam and sire, respectively, were mixed to produce each pairwise cross (i.e., family). Five dams and seven sires from Martin were crossed to produce 11 families in the Martin ✕ Martin (M ✕ M) within-reef offspring group (Figure 1C). Six dams and nine sires from Davies were crossed to produce 11 families in the Davies ✕ Davies (D ✕ D) within-reef offspring group. Six dams from Davies and five sires from Martin were crossed to produce six families in the Davies ✕ Martin (D ✕ M) between-reef offspring group. Six dams from Martin and seven sires from Davies were crossed to produce eight families in the Martin ✕ Davies (M ✕ D) between-reef offspring group. Fragments from each broodstock colony were collected for DNA analysis, preserved in ethanol, and stored at −20 °C until processing.

Larvae from the M ✕ M, D ✕ D, D ✕ M, and M ✕ D offspring groups (nine, seven, four and five families respectively) were sampled for DNA analyses at 4 – 8 days post fertilization to assess the effect of within- and between-reef crossing on the genetic diversity and differentiation of generated offspring groups. Individual larvae were collected for DNA analysis, preserved in ethanol, and stored at −20°C until processing (Figure 1C).

The effect of within- and between-reef crossing on the gene expression profiles under thermal stress was then assessed at the recruit stage. Larvae from the M ✕ M, D ✕ D, D ✕ M, and M ✕ D offspring groups (three, five, three and three families respectively) that had been maintained at 27.5°C were settled onto PVC slides using a synthetic peptide derived from *Hydra* sp. to induce settlement (Macadam et al., 2025; Takahashi et al., 1997). After successful settlement (≥ 1 polyp) and grow-out (30 days total; Figure 1C), recruits were transferred to six 50 L experimental tanks held at 27.5°C. Recruits were exposed to artificial lighting that emulated 08:00 sunrise and 16:00 sunset (two-hour ramp up and down respectively) with peak light levels exhibiting a photosynthetically active radiation (PAR) of approx. 70 μmol photons m⁻² s⁻¹. Recruits were tank acclimated and exposed to wild sediments assumed to contain Symbiodiniaceae symbionts collected from Wood Reef (northern GBR; −11.81181, 143.98162) for 39 days prior to the heat stress trial (Figure 1C; Macadam, et al., 2025). Three tanks were ramped at a rate of 0.5°C per hour to 32°C (heat stress treatment) and three tanks remained at 27.5°C (ambient treatment) with 0.8 L/min flow of 0.5 µM filtered water into the tanks resumed following attainment of treatment temperature. Recruits were held at these temperatures for the duration of the experiment, which concluded after 36 days of heat stress at 32°C. At the end of the experiment, all surviving singleton (i.e., discretely settled) recruits from the 27.5°C and 32°C treatments were preserved in RNAlater™ and stored at –20 °C until processing (Figure 1C).

### 2.2 Total RNA extraction

Total RNA was extracted from individual surviving singleton recruits from M ✕ M, D ✕ D, and M ✕ D (two families each) using the RNAqueous™ Micro Total RNA Isolation Kit (Invitrogen) following manufacturer instructions (see Supplementary Materials). Extracted total RNA underwent QA/QC using a Bioanalyzer 2100 (Agilent) with RNA 6000 Pico Kit chips (Agilent) by diluting a 1 µL aliquot of each sample by 1:10 – 1:50 in nuclease-free water, denaturing (70°C for 2 minutes), and loading 1 µL of denatured dilution onto individual RNA Pico chips (*n* = 11 samples per chip) along with manufacturer-supplied RNA-specific dye (containing internal standard) and RNA ladder to attain data regarding sample quantity (ng/µL) and integrity (RNA Integrity Number, RIN).

### 2.3 DNA and RNA sequencing

Ethanol preserved Martin and Davies broodstock samples (*n* = 9 and 11) as well as M ✕ M, D ✕ D, D ✕ M, and M ✕ D larvae (*n* = 44, 35, 20, and 25 respectively) were shipped frozen on dry ice in 96-well plates to Diversity Arrays Technologies (DArT; Canberra ACT Australia) for DNA extraction, QA/QC assessment, reduced representation genome sequencing, and SNP calling (Supplementary Table S1). Briefly, DArTseq utilizes Illumina HiSeq2500 next generation sequencing and endonuclease-facilitated genome reduction techniques to generate reduced representation genomic data with balanced site and genome coverage (Sansaloni et al., 2011).

Total RNA samples from individual singleton M ✕ M, D ✕ D, and M ✕ D recruits (*n* = 12, 14, and 14 respectively) were plated with wells containing 6 µL or 13 µL aliquots if concentration was ≥ 5 or < 5 ng/µL, respectively (Table S2). Plated RNA samples were sent frozen on dry ice to the Biomolecular Resources Facility (BRF) at Australian National University in Canberra ACT for library preparation using the NEBNext Ultra II kit (mRNA pull-down) followed by 150 base pair (bp) paired-end sequencing on an Illumina NovaSeq6000 using an entire S4 flowcell (300 cycles; see Supplementary Materials).

### 2.4 SNP analysis pipeline

The returned SNP sequence data were processed through the *DArT* pipeline using *de novo* clustering, which removed poor quality sequences, ensured reliable assignment of sequences to samples, and applied the *DArTsoft14* algorithm to identify SNPs at each locus across samples (Melville et al., 2017). The post-pipeline SNP data delivered by DArT for broodstock and larvae was further filtered using the package *dartR* v2 (Mijangos et al., 2022) in R (R Core Team, 2020) using seven sequential steps (see Supplementary Materials). Following SNP filtration, which retained all 20 broodstock and 120 of 124 larvae (two D ✕ D and two M ✕ D individuals removed), samples were subjected to principal coordinates analysis (PCoA) using Euclidian distances (Georges et al., 2023) to visualise the genetic structure within and among adult broodstock as well as within and among larvae from within- and between-reef offspring groups. Observed heterozygosity (H_o_) was calculated for individuals within the filtered SNP dataset then compared among the broodstock and offspring groups. Differences in group-level mean H_o_ were assessed using one-way analysis of variance (ANOVA) followed by Tukey’s HSD post-hoc tests (α = 0.05). Robustness of H_o_ estimates to sampling composition was evaluated using bootstrap resampling at both the individual and family levels (5,000 iterations), with mean H_o_ recalculated for each broodstock and offspring group in each replicate. Pairwise fixation indices (*F_ST_*) were estimated using Weir and Cockerham (1984) style allele-frequency estimators, which are robust to unequal numbers of sampled individuals among groups (Willing et al., 2012), with significance assessed using 5,000 bootstrap replicates across loci. Because offspring groups comprised related individuals, *F_ST_* values are interpreted as measures of genetic differentiation among structured cohorts rather than panmictic populations. The filtered SNP dataset was also used to calculate individual-level Euclidean genetic distances, which were summarized as within-group pairwise distances for each broodstock and offspring group and then assessed using one-way ANOVA followed by Tukey’s HSD post-hoc tests to determine differences in mean within-group genetic distance among groups.

### 2.5 RNAseq analysis pipeline

Raw 150 bp paired-end reads were processed through the *nf-core/rnaseq* pipeline v3.12.0 (Ewels et al., 2020) using *Nextflow* v23.10.0 (Di Tommaso et al., 2017). All software was run using *nf-core/rnaseq* v3.12.0 defaults except where specified. Reads processing with *nf-core/rnaseq* pipeline included adapter and quality trimming with *fastP* v0.23.4 (`min_trimmed_reads = 500`; Chen, 2023) before removal of human contaminated reads with *bbmap* v39.01 using Human genome hg38 as reference (Bushnell, 2014) and ribosomal RNA contaminated reads using *sortmerna* v4.3.4 (Kopylova et al., 2012). Retained reads were then mapped to an *A. tersa* (previously referred to as *A. hyacinthus* ‘neat’; Rassmussen et al., 2025; T. Bridge personal communication) reference genome (López-Nandam et al., 2023) with *STAR* v2.7.9a (Dobin et al., 2013) and the aligned reads quantified using *salmon* v1.10.1 (`min_mapped_reads = 0.001`; Patro et al., 2017).

RNAseq data was analysed in the within- and between-reef recruits from ambient (*n* = 20) and heat stress (*n* = 20) treatments (Figure 1C). To minimise the influence of low sequencing depth and optimise the robustness of the downstream analyses and inferences, samples with library size ≤ 2.5M were removed. Differential gene expression analyses were conducted in R v4.4.1 via R Studio v2024.09.0-375 using a conservative approach that considered the overlapping results of three discrete statistical tests performed using the following R packages for RNAseq analyses: *edgeR* (Chen et al., 2016; Robinson et al., 2010), *limma voom* (hereafter *limma*; Ritchie et al., 2015), and *DESeq2* (Love et al., 2014). All analyses were conducted using the same `model.matrix (∼ 0 + Group)` to compare group specific means as well as the same read count filtering stringency of counts per million (CPM) ≥ 10 for *n* ≥ 3 samples (i.e., expressed). To determine differentially expressed genes (DEGs), *edgeR* and *limma* analyses were conducted concurrently due to overlapping steps (e.g., trimmed mean of M values (TMM) normalization and `makeContrasts`) while *DESeq2* analysis was performed discretely.

For concurrent *edgeR* and *limma* analyses, the length scaled count matrix was imported, rounded to the nearest integer (for consistency with *DESeq2*), converted into a DGE object, and normalized using TMM (Robinson & Oshlack, 2010) before defining the contrast matrix, which specified the sample groupings for each contrast of interest (e.g., reef-level heat-stress response, frontloading, and interaction effects). For *edgeR* analysis, dispersions were estimated (`estimateDisp, robust = TRUE`) before a quasi-likelihood general linear model was fit (`glmQLFit, robust = TRUE`) and each contrast (`makeContrasts`) was assessed (by name) using the robust and error controlling quasi-likelihood F-test (`glmQLFTest`; Chen et al., 2016). Significant DEGs were determined using `topTags` as those with Benjamini-Hochberg adjusted *p* values < 0.05. For *limma* analysis, TMM-normalized read counts were transformed using `voomWithQualityWeights` (Liu et al., 2015, 2016) and fit to a linear model using `lmFit`. Moderated t-statistics were then generated via an empirical Bayes method using `eBayes` (Li et al., 2022) before significant DEGs (Benjamini-Hochberg adjusted *p* values < 0.05) were identified for each contrast (by coefficient number) using `topTable`. Lastly, *DESeq2* imported the same length scaled count matrix for each subset, rounded to the nearest integer, and converted into a `DESeqDataSet` object using `DESeqDataSetFromMatrix`. Following dispersion estimation and model fitting (*DESeq*), DEGs for the same contrasts defined in *edgeR*/*limma* contrast matrix were determined by Wald Test (Benjamini-Hochberg adjusted *p* values < 0.05) using the results command (`independentFiltering = FALSE`). Up- and down-regulated DEGs identified by *edgeR*, *limma*, and *DESeq2* (i.e., consensus DEGs) for each contrast were combined and appended with reference genome annotation (López-Nandam et al., 2023) using a custom R script. Tables reporting the top consensus DEGs for each contrast were generated by ranking all significant genes by the average of log_2_ fold-changes reported by *edgeR*, *limma*, and *DESeq2*. All consensus DEGs that lacked functional annotation (i.e., “uncharacterized protein” or “predicted protein”), top consensus DEGs with multiple identical functional annotations, and putative heat stress candidate genes were assessed for homology to annotated proteins by UniProt *BLASTP* searches of the UniProtKB reference proteomes and Swiss-Prot databases (v2.16.0+; Altschul et al., 1997; Schäffer et al., 2001). All unmapped reads were assessed against the NCBI core nucleotide database to identify potential origin by running *Kraken2* (Wood et al., 2019) using *nf-core/taxprofiler* v1.1.0 (Stamouli et al., 2023) with the following arguments: `--run_kraken2 --kraken2_save_readclassifications --run_profile_standardisation --run_krona`. Consensus DEGs with functional annotation that significantly responded to heat stress treatment in at least one offspring group were visualized as heatmaps (*pheatmap* v1.0.12; Kolde, 2018) that contained either filtered (log_2_ fold-change > 1) or unfiltered DEGs.

To identify the Gene Ontology (GO) terms and euKaryotic Orthologous Group (KOG) categories associated with each geneID based on protein homology, the *A. tersa* transcriptome-derived proteome (López-Nandam et al., 2023) was processed through *eggNOG-emapper* v2.1.12 (<u>eggnog-mapper.embl.de/</u>) using *eggnog5* and default parameters (Cantalapiedra et al., 2021; Huerta-Cepas et al., 2019). Each geneID in the count matrix was mapped to a KOG category (single or multiple letters) using a custom R script and then all geneIDs associated with each single-letter KOG category were tested for differences between ambient and heat stress treatments when offspring groups were considered together and separately using Mann Whitney U tests (Benjamini-Hochberg adjusted *p* value < 0.05) on CPM values (see above). For visualization, CPM values were averaged for each offspring group from each treatment, log_2_ transformed, row-wise (i.e., KOG-wise) z-score normalized, and plotted as hierarchical heat map with Euclidian distances for row clustering using *pheatmap* v1.0.12 (Kolde, 2018). The GO-slim generic subset (2025-06-01 release; v1.2; Aleksander et al., 2023; Ashburner et al., 2000) was loaded into a custom R script using `get_ontology` within *ontologyIndex* (Greene et al., 2017) then the `GOslim_term2gene` and `GOslim_term2name` objects were generated and included all terms from the *eggNOG-emapper* output file (*n* = 1,075,837) and the corresponding GO-slim generic terms (*n* = 133), respectively. Before gene set enrichment analysis (GSEA; Subramanian, et al., 2005) was run to identify enriched GO-slim generic terms, the count matrix was regularized log (rlog) transformed with consideration of the design (`blind = FALSE`) using *DESeq2* before log_2_ fold-changes were calculated in Excel by subtracting the ambient from the elevated treatment for each sample (Lier et al., 2022; Marx et al., 2022). GSEA was run on rlog log_2_ fold-change ranked gene lists using *fgsea* (Korotkevich et al., 2016) in *clusterProfiler* v4.12.6 (Wu et al., 2021) with GO-slim generic terms considered to be significantly enriched when Benjamini-Hochberg adjusted *p* value ≤ 0.01. All GO-slim generic terms that were significantly enriched in at least one offspring group were visualized as a heatmap generated by *pheatmap* v1.0.12 (Kolde, 2018).

## 3. Results

### 3.1 Genomic diversity and structure of larval and broodstock populations

The filtered SNP dataset contained 2,337 of the 25,535 loci identified by the *DArTsoft14* algorithm (9.2% of pre-filtration SNP dataset retained) for each of the 140 individuals (97.2% of pre-filtration SNP dataset retained; *n* = 120 larvae and *n* = 20 adult broodstock colonies).

The first two axes of a PCoA of this data explained 8.9% and 5.9% of the total genetic variation amongst all samples (Figure 2). Martin and Davies broodstock clustered together except for three Martin (M9, M13, and M14) and two Davies (D8 and D13) broodstock samples (Figure 2). Overall, the clustering of offspring groups was driven by familial relationships (i.e., whether they shared a sire and/or dam; Figure 2A) except for two within-reef offspring groups that did not cluster together despite sharing a dam (M14 ✕ M9 and M14 ✕ M3); however, these do sit halfway between their respective parental colonies (Figure 2B; Supplementary Materials). Notably, some families clustered together despite not sharing a parent (e.g., D16 ✕ M8 and D6 ✕ D15), which is potentially due to genetic similarities amongst the broodstock colonies.

**Figure 2.**
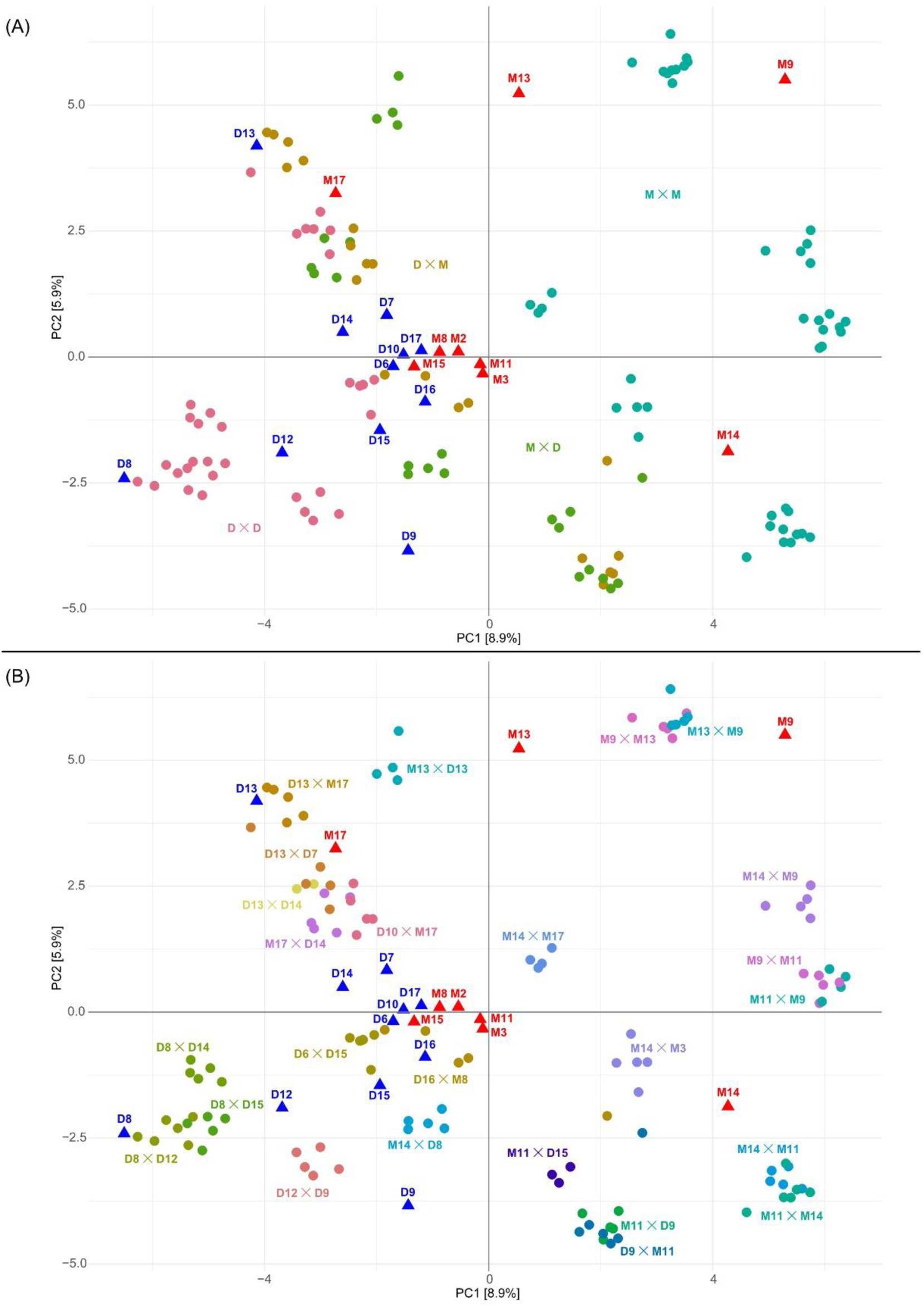
Principal coordinates analysis (PCoA) based on Euclidian distances of genome-wide single nucleotide polymorphisms (SNPs) depicting genetic relatedness among all *Acropora tersa* larvae from within-reef (Martin ✕ Martin, M ✕ M, *n* = 44; Davies ✕ Davies, D ✕ D, *n* = 33) and between-reef (Martin ✕ Davies, M ✕ D, *n* = 23; Davies ✕ Martin, D ✕ M, *n* = 20) crosses. Broodstock were sourced from of Martin (M, *n* = 9) and Davies (D, *n* = 11) Reefs. Circles and triangles represent individual larvae and adult colonies, respectively, with proximity indicating genetic relatedness. Colours denote broodstock plus offspring groups (A) or families (B). Labels list the dam colony followed by the sire colony (separated by ‘✕’).

When compared to each other, broodstock colonies (Martin versus Davies) and within-reef offspring groups (M ✕ M versus D ✕ D) exhibited the lowest and highest significant pairwise genetic differentiation, respectively, though differentiation remained modest overall (Table 1). M ✕ M and D ✕ D offspring groups exhibited the lowest genetic differentiation with their respective broodstock, respectively, as expected given shared parental origin. M ✕ M offspring exhibited lower differentiation with M ✕ D than D ✕ M offspring while D ✕ D offspring exhibited lower differentiation with M ✕ D than D ✕ M offspring. Consistent with their mixed parental composition, differentiation between D ✕ M and M ✕ D was lower than when either between-reef offspring group was compared to M ✕ M and D ✕ D (Table 1).

**Table 1.** Pairwise genetic differentiation and distance among *Acropora tersa* broodstock and offspring groups. Pairwise fixation indices (*F_ST_*; below diagonal) and Euclidean genetic distance comparisons (above diagonal) were calculated among broodstock from Martin (M) and Davies (D) reefs as well as larvae from within-reef (M ✕ M, D ✕ D) and between-reef (M ✕ D, D ✕ M) offspring groups using the filtered SNP dataset (2,337 loci). Bolded *F_ST_* and Euclidean distance values indicate statistical significance (*p* < 0.01 based on 5,000 bootstraps and Tukey HSD tests), respectively. Sample sizes (*n*) for each group are shown in parentheses. Labels list dam colony followed by sire colony (separated by ‘✕’).

| | M × M | M × D | D × D | D × M | D | M |
| --- | --- | --- | --- | --- | --- | --- |
| M × M ($n = 44$) | - | **1.73** | **1.70** | **1.22** | **3.86** | **3.35** |
| M × D ($n = 23$) | **0.055** | - | -0.02 | -0.51 | **2.14** | 1.63 |
| D × D ($n = 33$) | **0.131** | **0.04** | - | 0.49 | **2.16** | -1.65 |
| D × M ($n = 20$) | **0.09** | **0.025** | **0.072** | - | **2.64** | **-2.14** |
| D ($n = 11$) | **0.092** | **0.01** | -0.005 | **0.019** | - | 0.51 |
| M ($n = 9$) | **0.032** | **0.015** | **0.062** | **0.024** | **0.008** | - |

Euclidian genetic distances revealed the same pairwise comparison patterns as *F_ST_* (Table 1; Supplementary Figure S1A). Further, genetic distances within each broodstock and offspring group revealed one or three relatedness peaks (e.g., non-, half-, or full-sibling) with Martin and Davies broodstock being exclusively comprised of non-sibling colonies and each family comprised of full-sibling individuals; however, relatedness structure within offspring groups was heterogeneous, most notably an enrichment in non-sibling M ✕ D individuals (Figure S1B).

Filtered SNPs revealed an overall H_o_ of 0.20 ± 0.03. M ✕ D offspring exhibited significantly higher H_o_ than M broodstock (Tukey HSD *p* = 0.0292) whereas no other pairwise comparisons were significant (Tukey HSD *p* > 0.07; Figure 3). H_o_ estimates were robust to variation in the number of families and larvae that contributed to each offspring group, as indicated by bootstrap resampling (Figure S2; Table S3).

**Figure 3.**
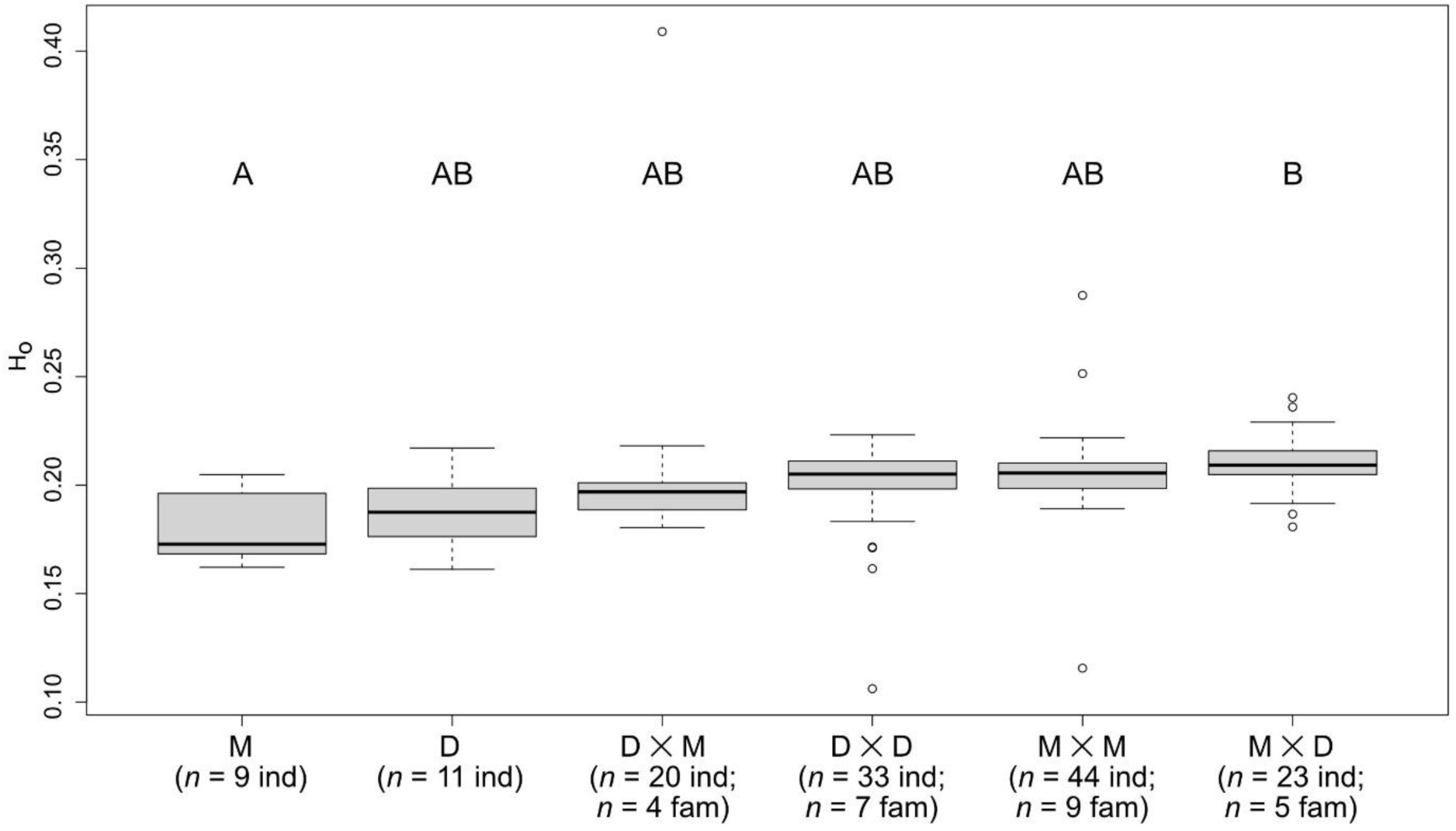
Boxplot of the observed heterozygosity (H_o_) among *Acropora tersa* larvae generated by within-reef (Martin ✕ Martin, M ✕ M and Davies ✕ Davies, D ✕ D; *n* = 44 and *n* = 33) and between-reef (Martin ✕ Davies, M ✕ D and Davies ✕ Martin, D ✕ M; *n* = 23 and *n* = 20) crosses of broodstock from Martin and Davies reefs (M and D; *n* = 9 and *n* = 11), respectively. A significant difference in H_o_ was observed only between Martin reef broodstock and M ✕ D offspring group as denoted by the unique letters above boxes (Tukey HSD *p* = 0.0292). H_o_ estimates were robust to variation in the number of families and larvae that contributed to each offspring group (Supplementary Figure S2; Supplementary Table S3). Boxplots are ordered by median H_o_ while statistical comparisons are based on mean H_o_ values. Labels list dam colony followed by sire colony (separated by ‘✕’) as well as the number of individual (ind) larvae that came from the number of generated families (fam).

### 3.2 Recruit transcriptional responses to heat stress

Mapping of cleaned paired end reads (1,271,258,532 total for 40 samples) to the *A. tersa* reference genome (27,110 genes) yielded library sizes of 2,821,139 – 41,521,050 read pair counts (mapping rate 90.9 ± 0.5% including unique and multi-mapped read pairs) after removal of one sample due to library size < 2.5M. KOG mapping, consensus DEG determination, and GSEA analysis were conducted using the retained 12,531 (47.7%) genes per sample after non-and lowly-expressed genes were removed by CPM filtration, of which 7,591 (60.6%) and 7,470 (59.6%) had an associated KOG category and GO-slim generic terms, respectively. Assessment of all unmapped reads did not identify any with annotation to Symbiodiniaceae, which aligns with the observation that recruit pigmentation remained pale to the naked eye throughout the experiment (Figure S3); therefore, all results pertain to host only.

Based on previous evidence for strong maternal and population-specific influences on early life stage thermal tolerance, we expected recruit transcriptional responses to differ primarily between within-reef and between-reef crosses, with stronger divergence among maternal lineages (i.e. D ✕ D vs M ✕ M), and intermediate responses in mixed crosses (M ✕ D). Contrary to this expectation, recruit transcriptional responses did not exhibit clear separation between D ✕ D, M ✕ M, and M ✕ D offspring groups. Although overlap of differentially expressed genes among all offspring groups was limited, the dominant transcriptional responses converged on similar stress-response pathways and functional categories rather than distinct cross-specific expression programs.

#### 3.2.1 KOG functional categories

Across all recruit offspring groups, the majority of CPM values for the 7,591 expressed genes that mapped to a KOG category were associated with the 24 non-overlapping and non-predictive single letter categories (93.7%; Figure 4).

**Figure 4.**
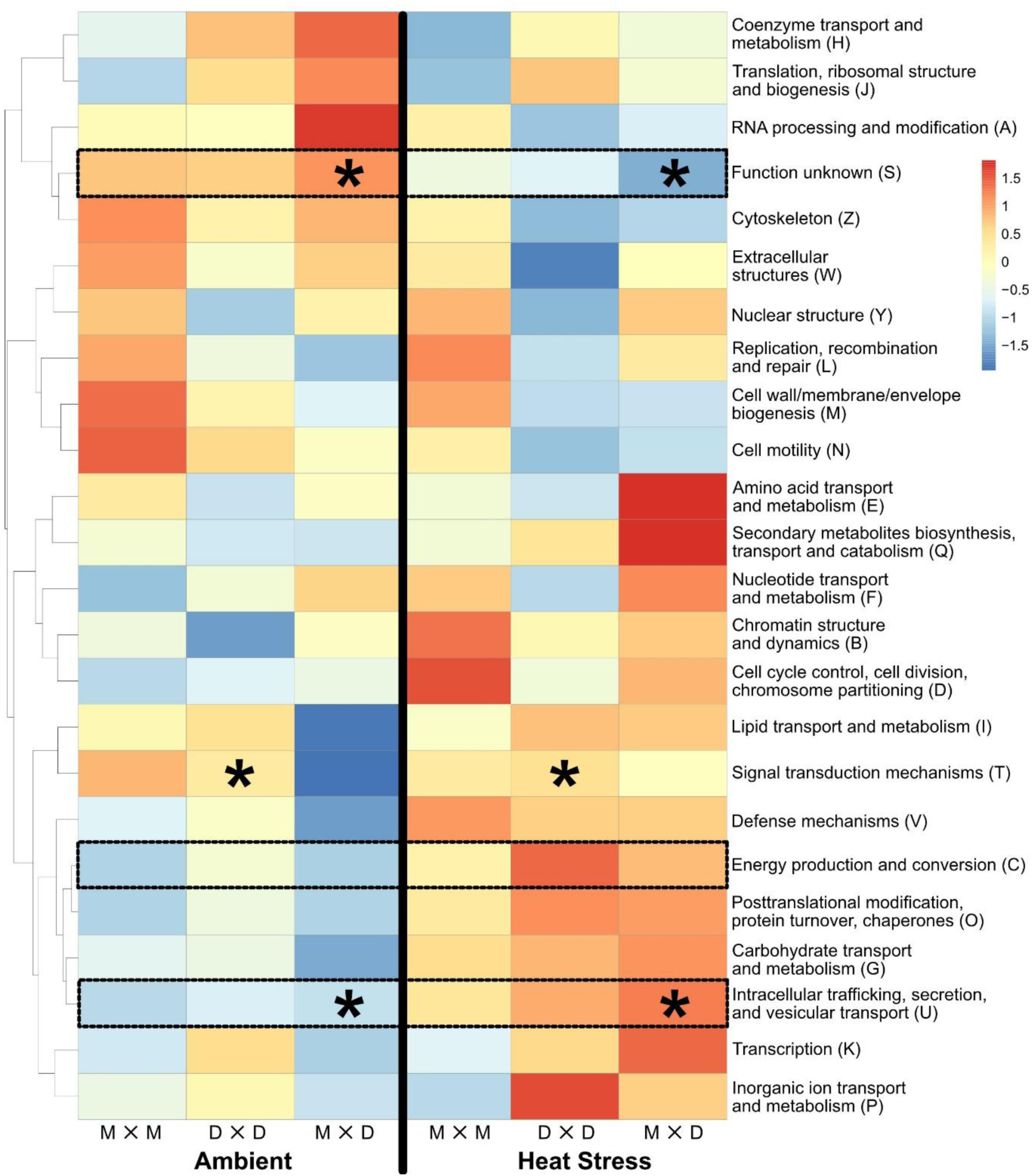
Heat map of euKaryotic Orthologous Group (KOG) category functional profiles of *Acropora tersa* under ambient (27.5°C) and heat stress (32°C) conditions. Colours represent row-wise (i.e., KOG-wise) z-score normalized differences of counts per million values for the 7,591 (of 12,531) genes that exhibited protein-based homology with KOG categories for Martin ✕ Martin (M ✕ M), Davies ✕ Davies (D ✕ D), and Martin ✕ Davies (M ✕ D) recruits (*n* = 12, 14, and 14 averaged individuals respectively). The majority of CPM values (93.7%) were captured by these 24 non-overlapping and non-predicted single-letter categories, demonstrating comprehensive transcriptome coverage. For each KOG category, box outlines indicate a significant difference between treatments across offspring groups while asterisks indicate a significant difference between treatments within an offspring group (Mann Whitney U Test adjusted p value < 0.05). Offspring group labels list dam colony followed by sire colony (separated by ‘✕’).

Under ambient conditions, expression levels varied subtly within and among recruit offspring groups, including elevated cell wall/membrane/envelope biogenesis (M) and cell motility (N) genes in M ✕ M recruits, decreased nuclear structure (Y) and chromatin structure and dynamics (B) genes in D ✕ D recruits, and elevated RNA processing and modification (A) alongside reduced lipid transport and metabolism (I), signal transduction mechanisms (T), defence mechanisms (V), and carbohydrate transport and metabolism (G) genes in M ✕ D recruits.

Across the 23 annotated categories, heat stress was associated with coordinated induction of stress-related and homeostatic functional categories alongside suppression of growth-, signalling-, and structural-related processes. More specifically, responses to heat stress were conserved across offspring groups and characterised by consistent elevation in defence mechanisms (V), energy production and conversion (C), posttranslational modification, protein turnover, and chaperones (O), carbohydrate transport and metabolism (G), intracellular trafficking, secretion and vesicular transport (U), and transcription (K) genes. Although magnitude of response varied modestly among crosses, with M ✕ M recruits generally exhibiting milder responses and D ✕ D recruits showing stronger responses, these differences did not translate into distinct expression profiles among cross types. Consistent with this pattern, D ✕ D recruits showed stronger responses in extracellular structure (W), energy production and conversion (C), and inorganic ion transport and metabolism (P) while M ✕ D recruits exhibited relatively stronger responses in amino acid transport and metabolism (E) and secondary metabolite biosynthesis, transport, and catabolism (Q).

When offspring groups were considered together (Figure 4), CPM values differed significantly between ambient and heat stress treatment conditions for genes associated with unknown function (Mann Whitney U adjusted p < 0.01), energy production and conversion (Mann Whitney U adjusted p < 0.05), and intracellular trafficking, secretion, and vesicular transport (Mann Whitney U adjusted p < 0.01). When offspring groups were analysed separately, significant treatment effects were limited to specific functional categories, including signal transduction mechanisms (T) genes in D ✕ D recruits (adjusted *p* < 0.01) as well as unknown function (S) and intracellular trafficking, secretion, and vesicular transport (U) genes in M ✕ D recruits (adjusted *p* < 0.0001 and *p* < 0.01 respectively).

#### 3.2.2 Differential expression

Consistent with the broadly conserved functional stress responses identified by KOG analyses, gene-level differential expression patterns were examined to assess how individual transcripts contributed to these shared responses. All significantly up- and down-regulated consensus DEGs observed under heat stress relative to ambient conditions for the within- and between-reef recruit groups (i.e., consensus heat stress DEGs; Table 2) were used to characterise transcriptional responses. While differential expression was assessed across all genes, results are summarised for consensus heat stress DEGs with absolute log₂ fold-changes (FC) ≥ 1 (i.e., ≥ 2-fold) and functional annotations to aid biological interpretation (Figure 5; Figures S4 and S5; Table S4). Across offspring groups exposed to heat stress relative to ambient, M ✕ M recruits (*n* = 6 per treatment) displayed a relatively modest transcriptional response (28 up- and 30 down-regulated DEGs; log_2_ FC ∼0.5 – 1.9 and −0.4 – −2.0; Supplementary File S1), D ✕ D recruits (*n* = 7 per treatment) showed a markedly larger response (252 up- and 128 down-regulated DEGs; log_2_ FC ∼0.3 – 5.6 and −0.4 – −2.5; File S2), and M ✕ D recruits (*n* = 7 per treatment) exhibited an intermediate response (44 up- and 45 down-regulated DEGs; log_2_ FC ∼0.5 – 3.7 and −0.4 – −2.1; File S3). Despite these differences in DEG magnitude and number, functional responses were broadly similar across offspring groups. Consistent with KOG (Section 3.2.1) and GSEA (Section 3.2.3) results, many up-regulated genes had annotations suggestive of cellular defence, protein homeostasis, metabolic reprogramming, immune signalling, and cellular architecture whereas many down-regulated genes had annotations suggestive of cell signalling and development, cytoskeletal organisation, gene regulation, and metabolic processes. UniProt *BLASTP* searches confirmed that up- and down-regulated consensus heat stress DEGs with identical annotations in D ✕ D recruits exhibited high sequence homology to the same annotated *Acropora cervicornis* proteins (Table S5). Taken together, these gene-level expression patterns mirror the conserved stress-related functional shifts observed in KOG analyses, including coordinated directional responses characterised by induction of defence, protein homeostasis, and metabolic processes alongside repression of growth- and signalling-related functions.

**Figure 5.**
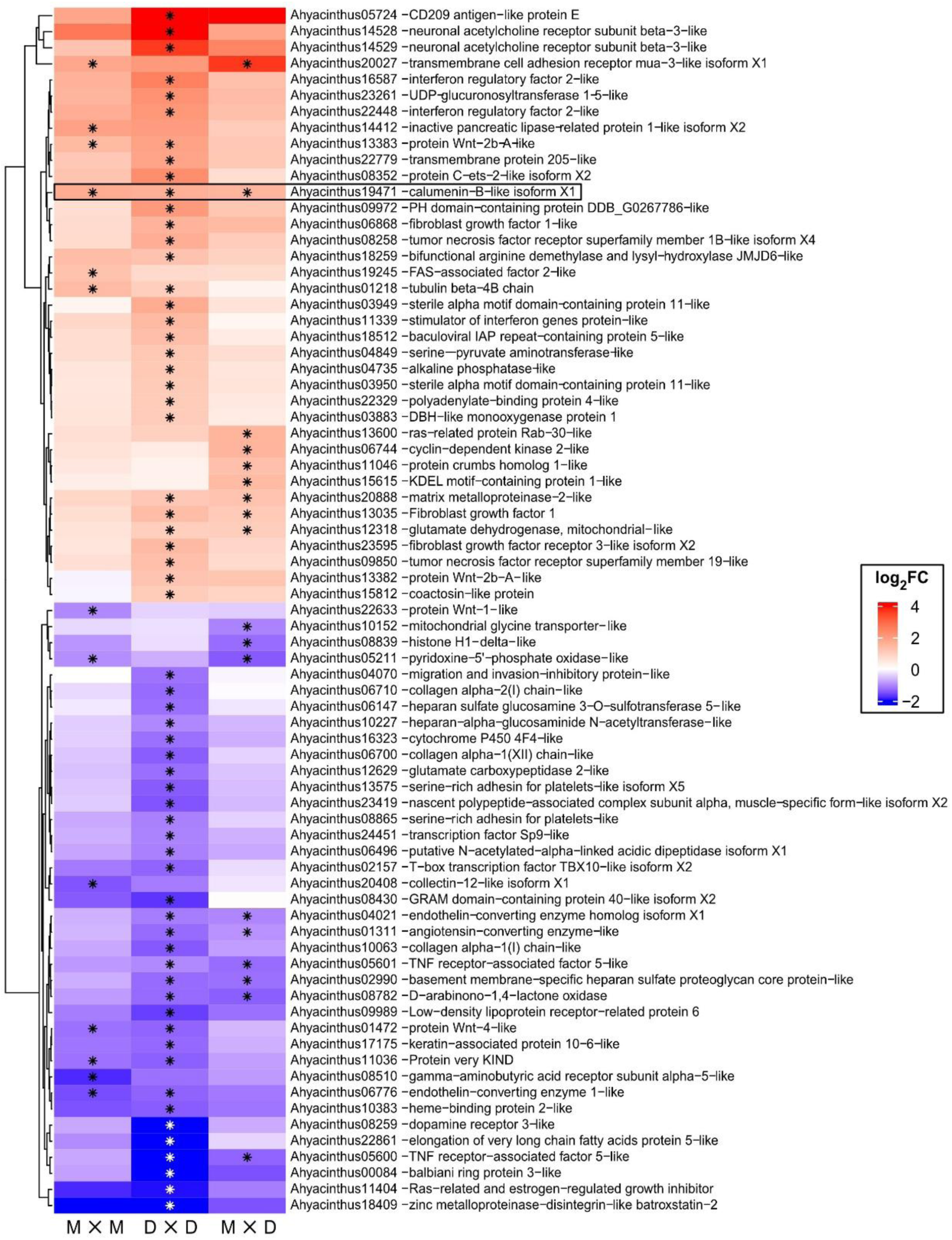
Heat map of differentially expressed genes of *Acropora tersa* recruits under heat stress (32°C) relative to ambient (27.5°C) conditions. Rows represent significant differentially expressed genes (DEGs) with log_2_ fold-change ≥ 1 or ≤ −1 and functional annotations in Martin ✕ Martin (M ✕ M), Davies ✕ Davies (D ✕ D), and Martin ✕ Davies (M ✕ D) recruit offspring groups (*n* = 12, 14, and 14 averaged individuals respectively; see Supplementary Figure S4 for all DEGs with functional annotations). Asterisks denote DEGs that were identified as significant by *edgeR*, *limma voom*, and *DESeq2* (i.e., consensus heat stress DEGs; Benjamini-Hochberg adjusted *p* value < 0.05). Box outline highlights one of the seven genotype-independent consensus heat stress DEGs with log_2_ fold-change ≥ 1 (see Supplementary Figure S4 for all). Shades of red and blue represent the magnitude of up- and down-regulation, respectively. Offspring group labels list dam colony followed by sire colony (separated by ‘✕’).

**Table 2.** Summary of significant up- and down-regulated differentially expressed genes (DEGs) identified by *edgeR*, *limma voom*, and *DESeq2* in *Acropora tersa* recruits under heat stress relative to ambient conditions. Consensus DEGs are those identified as significant in all three differential expression analyses. DEG counts are shown for within-reef (Martin ✕ Martin, M ✕ M and Davies ✕ Davies, D ✕ D) and between-reef (Martin ✕ Davies, M ✕ D) offspring groups. Offspring groups list dam colony followed by sire colony (separated by ‘✕’).

| Cross | Effect | *edgeR* |  | *limma voom* |  | *DESeq2* |  | Consensus |  |
| --- | --- | --- | --- | --- | --- | --- | --- | --- | --- |
|  |  | Up | Down | Up | Down | Up | Down | Up | Down |
| M × M | Main | 42 | 36 | 78 | 72 | 56 | 52 | 28 | 30 |
| D × D | Main | 305 | 195 | 419 | 218 | 360 | 303 | 252 | 128 |
| M × D | Main | 102 | 77 | 51 | 55 | 202 | 152 | 44 | 45 |
| M × M vs<br>D × D | Interaction | 0 | 0 | 0 | 0 | 7 | 8 | 0 | 0 |
| M × M vs<br>M × D | Interaction | 2 | 0 | 0 | 0 | 8 | 7 | 1<sup>^</sup> | 0 |
| D × D vs<br>M × D | Interaction | 1 | 0 | 0 | 0 | 28 | 20 | 1<sup>^</sup> | 0 |
| M × D vs<br>M × M | Frontloaded | 18 | | 1 | | 49 | | 1 | |
| M × D vs<br>D × D | Frontloaded | 34 | | 15 | | 130 | | 9 | |
| M × M vs<br>D × D | Frontloaded | 48 | | 62 | | 80 | | 18 | |

<sup>^</sup>Interaction effect identified by *edgeR* and *DESeq2*.

To further resolve how gene-level transcriptional responses were distributed across offspring groups, consensus heat stress DEGs were examined for the extent of overlap among within- and between-reef recruits under heat stress relative to ambient conditions (Figure 5; Figures S4 and S5; Table S6), which identified: (1) six (*calumenin-B-like isoform X1, heat shock protein HSP 90-beta-like*, *calreticulin-like isoform X1*, *endoplasmin-like isoform X1*, *protein disulfide-isomerase A4-like*, and *formin-binding protein 4-like*) and one (*patched domain-containing protein 3*) among all within- and between-reef offspring groups (File S4); (2) 21 and seven between M ✕ M and D ✕ D (File S5), (3) seven and seven between M ✕ M and M ✕ D (File S6), and (4) 24 and 14 between D ✕ D and M ✕ D (File S7), respectively. All other significantly up- and down-regulated consensus heat stress DEGs with functional annotations were unique to each offspring group, which included four and 10 for M ✕ M (File S8), 187 and 96 for D ✕ D (File S9), and 16 and 20 for M ✕ D offspring groups (File S10), respectively. Notably, the D ✕ D offspring group (D8 ✕ D14 and D16 ✕ D17) consisted of non-sibling families whereas M ✕ M (M14 ✕ M3 and M9 ✕ M14) and M ✕ D (M15 ✕ D12 and M2 ✕ D12) offspring groups each consisted of half-sibling families. Of the nine consensus heat stress DEGs with non-functional annotations that were either common across all offspring groups or unique to between-reef recruits, UniProt *BLASTP* identified putative annotations for two of four or two of five, respectively (Tables S7 and S8).

When within- and between-reef offspring groups were assessed for consensus heat stress DEGs with significant interaction effects (i.e., where the effect of temperature treatment on differential gene expression was dependent on offspring group) none were identified; however, both *edgeR* and *DESeq2* identified one up-regulated DEG involved in transport and cellular homeostasis (*cationic amino acid transporter 2-like*) that exhibited a significant interaction effect between M ✕ M and M ✕ D as well as D ✕ D and M ✕ D (Table 2).

Comparisons among recruits subjected to ambient treatment conditions (Figure 1C) were also conducted to identify potentially frontloaded (i.e., higher baseline expression) genes. Significantly increased expression was observed for one and nine genes in M ✕ D recruits relative to M ✕ M and M ✕ D recruits (Files S11 and S12), respectively, as well as 18 genes in M ✕ M recruits relative to D ✕ D recruits (File S13). Considering all potentially frontloaded genes, three (*soma ferritin-like isoform X1*, *TNF receptor-associated factor 3-like isoform X1*, and *serine-rich adhesin for platelets-like*) exhibited significantly increased expression in both M ✕ D and M ✕ M recruits relative to D ✕ D recruits.

#### 3.2.3 GSEA using GO-slim

To evaluate whether the conserved functional responses to heat stress relative to ambient conditions that were identified by KOG analyses and directional gene-level patterns observed in DEG analyses were also reflected at the pathway enrichment level, GSEA was performed using GO-slim generic terms (Figure 6). M ✕ M recruits exhibited significant induction of genes associated with intracellular protein transport and protein folding alongside significant reduction of gene sets linked to cell adhesion, extracellular matrix organisation, extracellular matrix components, molecular transducer activity, and receptor–ligand activity (adjusted *p* < 0.01; File S14). D ✕ D recruits exhibited a broader suite of enriched GO-slim terms that included significant induction of genes associated with defence response to other organisms, generation of precursor metabolites and energy, intracellular protein transport, mitochondrion organisation, cytoplasmic translation, protein folding, and oxidoreductase activity alongside significant reduction of genes associated with establishment or maintenance of cell polarity, nervous system processes, and extracellular matrix components (adjusted *p* < 0.01; File S15). M ✕ D recruits exhibited significant induction of genes associated with protein folding and GTPase activity alongside significant reduction in genes associated with cytoplasmic translation, ribosomal components, molecular transducer activity, and structural molecular activity (adjusted *p* < 0.01; File S16).

**Figure 6.**
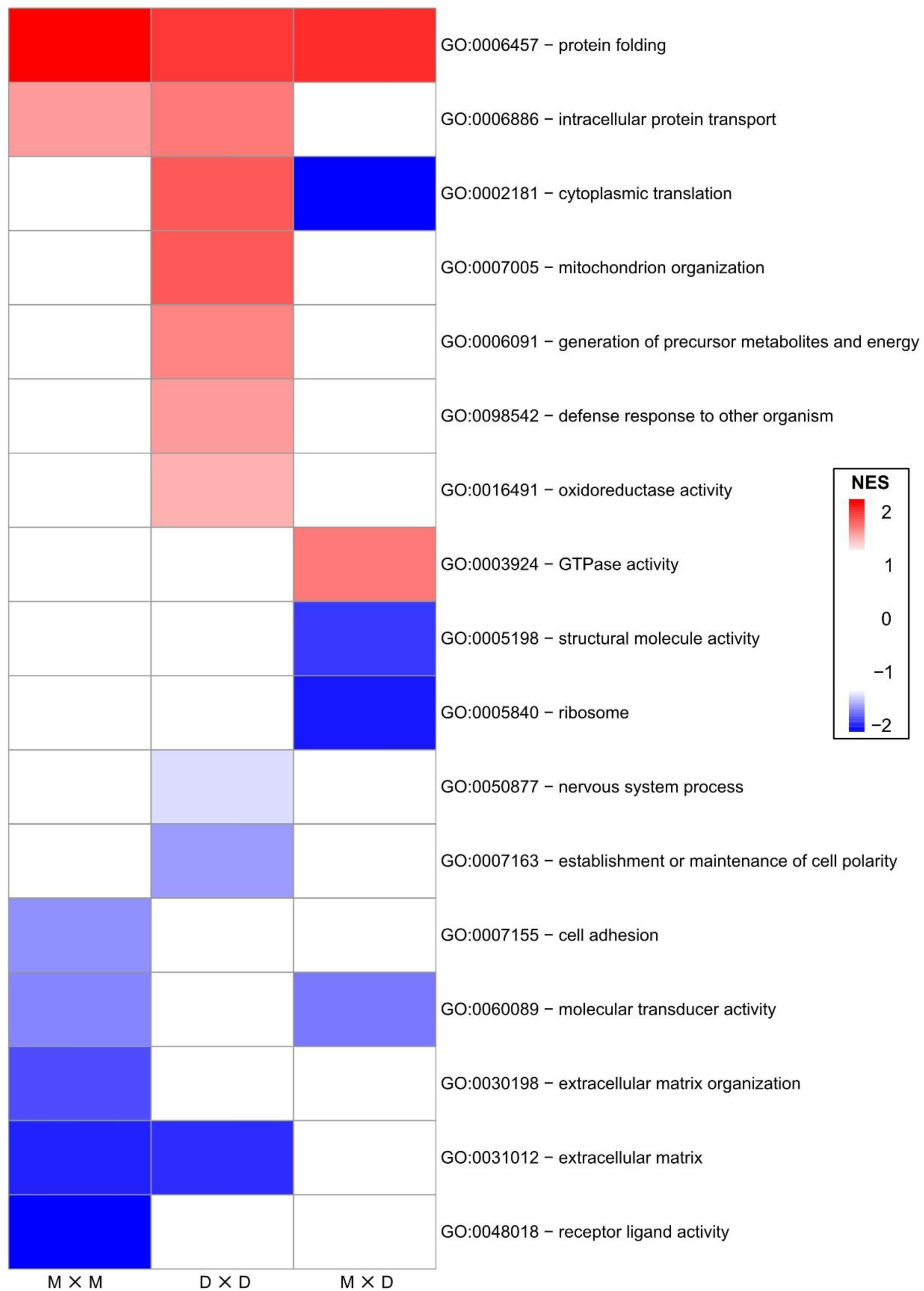
Heat map of gene ontology (GO)-slim generic subset term enrichment of *Acropora tersa* recruits under heat stress (32°C) relative to ambient (27.5°C) conditions. Rows represent significantly enriched GO-slim terms identified in at least one offspring group by gene set enrichment analysis (GSEA) conducted on the regularized log transformed counts per million (CPM) filtered dataset (*n* = 12,531) for Martin ✕ Martin (M ✕ M), Davies ✕ Davies (D ✕ D), and Martin ✕ Davies (M ✕ D) recruits (*n* = 12, 14, and 14 averaged individuals respectively). Colours represent normalized enrichment score (NES) values (≥ 1.3 or ≤ −1.3). Offspring group labels list dam colony followed by sire colony (separated by ‘✕’).

Despite differences in enrichment strength and the breadth of significant gene sets, several pathway-level responses were shared across offspring groups under heat stress: (1) protein folding associated genes were induced in all offspring groups (File S17); (2) intracellular protein transport associated genes were induced while extracellular matrix genes were reduced in both within-reef families (File S18), (3) molecular transducer activity associated genes were reduced in M ✕ M and M ✕ D recruits (File S19), and (4) cytoplasmic translation associated genes were induced in D ✕ D and reduced in M ✕ D recruits (File S20). Under ambient conditions (i.e., frontloading comparisons), GSEA identified enrichment of GO-slim terms related to cytoplasmic translation, intracellular protein transport, ribosome biogenesis, and ribosomal components in both M ✕ D and D ✕ D recruits relative to M ✕ M (Figure S6; Files S21 and S22 respectively). All remaining induced GO-slim terms under ambient conditions were unique to M ✕ M, D ✕ D, and M ✕ D recruits and limited in number (*n* = 0, 4, and 1; File S23, S24, and S25), respectively.

Variation among recruit offspring groups was driven largely by differences in enrichment strength and scope, not by engagement of distinct pathways. Therefore, GSEA results reinforce KOG and DEG analyses by demonstrating that heat stress responses across recruit offspring groups converge on a shared set of core stress response and homeostatic pathways (particularly protein folding, intracellular transport, and metabolic reprogramming), which are consistent with the Type A General Coral Stress Response (Dixon, et al., 2020).

## 4. Discussion

### Genetic diversity and differentiation

SNP-based genotyping provided valuable insights into the genetic diversity and differentiation of wild broodstock and their within- and between-reef offspring. Given the low genetic differentiation between Martin and Davies broodstock (Table 1), which is consistent with the moderate-to-high levels of gene flow documented along the GBR (Lukoschek, et al., 2016; Meziere, et al., 2025), the generally consistent levels of Hₒ across adult colonies and larval offspring groups were unsurprising. However, one between-reef offspring group (M ✕ D; four Martin and five Davies parents) exhibited significantly higher Hₒ than Martin broodstock whereas the reciprocal cross (D ✕ M; four Davies and three Martin parents) did not despite sharing one Martin and two Davies parents with M ✕ D, which suggests that broodstock-to-offspring shifts in Hₒ can vary non-uniformly among parental combinations (e.g., Quigley, et al., 2020b; Dallmeyer-Drennen et al., 2026). Notably, the significantly elevated Hₒ in M ✕ D did not correspond to enhanced larval survival under experimental heat stress (Macadam et al., 2025). Consistent with previous findings, between-reef larvae tended to exhibit higher mean Hₒ than within-reef offspring groups (Table S3); however, these differences were not statistically significant (Figure 3). This indicates that variation in offspring Hₒ is primarily driven by broodstock genetic composition rather than provenance alone. Given the importance of maintaining or increasing genetic diversity in corals for restoration purposes (Doropoulos et al., 2019; van Oppen et al., 2015), these findings highlight the need to understand genetic differentiation among targeted broodstock as this influences the genetic diversity and thus adaptive capacity of offspring generated through thermal history provenancing.

### Overall gene expression patterns under heat stress are conserved

Overall, consensus heat stress DEGs and GO-slim enrichment aligned with a conserved environmental stress response, characterized by the up-regulation of genes associated with DNA damage repair, protein folding and degradation, cell wall modification, intracellular signalling, vacuolar and mitochondrial functions, detoxification, carbohydrate and fatty acid metabolism, metabolite transport, and autophagy alongside the down-regulation of genes associated with growth-related processes (Gasch et al., 2000). This gene expression patterning is common across multiple *Acropora* species and referred to as the Type A General Coral Stress Response, with the Type B response being broadly opposite and generally observed following exposure to less intense stress (Dixon et al., 2020). The consistent expression of a canonical Type A response across all recruit offspring groups indicates that thermal history provenancing did not alter the core heat stress response, but differences in DEG number and effect size suggest variation in the overall scale of the transcriptional response among crosses (Table 2). A similar pattern of largely conserved transcriptional responses across offspring groups has been reported in *A. tenuis* recruits following heat stress exposure (Strader & Quigley, 2022). Although selective breeding based on thermal history can enhance heat tolerance phenotypes in some contexts (Howells et al., 2021; Quigley, et al., 2021; Kler Lago et al., 2025), the recruits generated here exhibited both conserved transcriptional responses (Figures 3 – 5) and physiological phenotypes (Macadam et al., 2025). Together, this study indicates that thermal history provenancing-informed selective breeding in low-differentiation systems may not produce clearly divergent molecular or phenotypic responses among offspring groups despite variation in gene expression magnitude.

Phenotypic assessment (survival, growth, and bleaching, i.e., paling) of the same recruit offspring groups showed no significant differences among within- and between-reef recruits under heat stress relative to ambient conditions; however, survival was uniformly high (> 65%) across all offspring groups indicating that the applied heat stress elicited a moderate rather than severe response (Macadam et al., 2025). The largely overlapping transcriptional responses observed across recruit offspring groups observed in this study are consistent with these findings. Although a small number of consensus heat stress DEGs were unique to each recruit offspring group (Figure S5; Files S8 – S10), the lack of significant phenotypic divergence suggests that these differences likely reflect fine-scale regulatory variation that may not translate into detectable differences in organismal performance under the moderate heat stress conditions tested here.

The differing MMM between Martin and Davies reefs (28.63 °C and 28.39 °C, respectively; Figure 1B) means that exposure to 32 °C for 36 days imposed approximately one additional DHW of cumulative thermal stress on D ✕ D compared to M ✕ M recruits (13.4 versus 12.2 DHW). This difference in cumulative thermal stress may contribute to the greater number and effect size of consensus heat stress DEGs observed in D ✕ D relative to M ✕ M recruits (Table 2; Figure S4). DHWs of stress accumulated by the between-reef offspring cannot be calculated since their parents come from reefs with different MMMs; however, M ✕ D recruits exhibited an intermediate number of DEGs compared to the within-reef offspring groups. Moreover, the *10 kDa heat shock protein* showed ≈ 2-fold up-regulation in D ✕ D and M ✕ D recruits but not in M ✕ M recruits (Figure S4; File S7). Up-regulation of this gene has been associated with heat stress and bleaching responses in *Millepora complanata* where it was significantly elevated in bleached relative to unbleached colonies (Elizárraga et al., 2023), which further suggests that D ✕ D and M ✕ D recruits may have experienced a relatively stronger heat stress exposure than M ✕ M recruits. While genotype can contribute substantially to transcriptional variance under environmental stress (Chille et al., 2024), the larger number and magnitude of consensus heat stress DEGs in the D ✕ D offspring group, which comprises non-sibling families, may partly reflect greater transcriptional variation associated with differences in genetic background among families (Meyer et al., 2011). However, the broadly conserved pathway-level responses observed here indicate that heat stress exposure was likely the primary driver of transcriptional responses, with genetic background and relatedness within offspring groups contributing less substantially.

Alternatively, the broadly similar transcriptional responses observed in M ✕ M and M ✕ D recruits raise the possibility of maternally influenced regulatory effects on gene expression. Evidence for maternal effects in corals has previously been reported at early life stages, including maternal influences on fertilisation success (van Oppen et al., 2014) as well as the constitutive presence of maternal transcripts involved in methylation and transcriptional regulation across life stages in *Montipora capitata* (Chille et al., 2021). In addition, long-lasting maternal contributions to gene expression have been observed in first-generation interspecific hybrids of *A. tenuis* and *A. loripes* (Chan et al., 2021). However, despite the similarity in differential gene expression in M ✕ D and M ✕ M recruits, the between-reef recruits exhibited a unique heat stress response when considering gene ontology, most notably the opposite response of genes associated with cytoplasmic translation relative to D ✕ D (Figure 6), which provides additional evidence that a different heat stress intensity was experienced (Dixon et al., 2020). Moreover, genes associated with structural molecule activity and mRNA metabolic process were only frontloaded in M ✕ D relative to M ✕ M whereas genes associated with protein folding were only frontloaded in D ✕ D relative to M ✕ M (Figure S6). This suggests that cross-specific regulatory differences that do not manifest as phenotypic differences may arise, which could be due to the generation of a novel epigenetic profile when crossing broodstock based on thermal history provenance irrespective of genetic distinctiveness given the response of the methylome to seasonal variability (Rodríguez-Casariego et al., 2020). Whole genome bisulfite sequencing demonstrated that epigenetic inheritance does occur in hard corals (Liew et al., 2020) but on a per-locus basis (Peterson et al., 2024). Taken together, future coral breeding initiatives based on thermal history provenancing should consider undertaking RNAseq in conjunction with ecological epigenetics tools (e.g., Dixon and Matz, 2021) to permit interpretation of the distinct transcriptional response in offspring based on their methylation signatures (*sensu* Dixon et al., 2014).

### Candidate thermal stress response markers

Gene expression in early (i.e., pre-adult) life history stages of corals has been relatively understudied (Walker et al., 2024); therefore, putative heat stress response candidate genes are limited (e.g., Ruggeri et al., 2023; Strader & Quigley, 2022). Here we consistently observed six and one significantly up- and down-regulated consensus heat stress DEGs across all recruit offspring groups (i.e., genotype-independent), respectively (Tables S6 and S8). *Calumenin-B-like isoform X1* (*calumenin-B*) likely encodes a calcium-binding protein involved in calcium homeostasis, apoptosis, and host-symbiont signalling (Ganot et al., 2011; Honoré, 2009; Orrenius et al., 2003) that has been associated with bleaching sensitivity in adult corals depending on pre-conditioning (Bellantuono et al., 2012). The observed up-regulation of *calumenin-B* was consistent with the up-regulation of *calumenin-A* in *P. astreoides* from inshore and offshore populations subjected to similar heat stress conditions (e.g., recruits exposed to 30.9 ± 1.1°C for 16 days instead of 32°C for 36 days; Ruggeri et al., 2023); however, *calumenin* was not reported as up-regulated in *A. tenuis* recruits exposed to 32°C for 56 days (Strader & Quigley, 2022). *Heat shock protein HSP 90-beta-like* (*HSP90*) likely encodes the molecular chaperone HSP90, which has been associated with rapid heat stress response in corals (Louis et al., 2017). Elevated expression of *HSP90* was observed in adult *A. palmata* fragments following one day but not two days of 32°C heat stress exposure (DeSalvo et al., 2010), adult *Montastraea cavernosa* fragments following 3 days but not 20 days of 33°C heat stress exposure (Skutnik et al., 2020), and juvenile *A. tenuis* following approx. 1 day of 32°C heat stress exposure (Yuyama et al., 2012). As such, this study is, to our knowledge, the first observation of elevated *HSP90* expression in coral recruits following prolonged exposure to moderate heat stress conditions (36 days at 32°C). *Calreticulin-like isoform X1* (*calreticulin*) likely encodes an endoplasmic reticulum resident calcium-binding protein involved in protein folding regulation along with heat shock proteins. *Calreticulin* was up-regulated in *A. tenuis* juveniles following approx. 1 day exposure to 32°C heat stress (Yuyama et al., 2012), which, taken together with the up-regulation of *calreticulin* and *HSP90* observed here, supports the functioning of *calreticulin* as a molecular chaperone (Henle et al., 1998). *Protein disulfide-isomerase A4-like* (*PDIA4*) likely encodes an endoplasmic reticulum resident enzyme involved in protein folding (Bowley et al., 2017; Xiong et al., 2020) and up-regulation of *PDIA4* and *calreticulin* was also observed in the marine invertebrate *Ruditapes philippinarum* following 1 day of 30°C heat stress exposure (Jing et al., 2023); however, a protein disulfide isomerase transcript was observed to be down-regulated in adult fragments of the offshore-sourced star coral *Orbicella faveolata* subjected to 5 days of 33°C heat stress (Aguilar et al., 2024). Given the inconsistent observations, *PDIA4* requires further investigation in corals, which is supported by the recent meta-analysis that identified two protein disulfide isomerase family A members (*PDIA3* and *PDIA6*) as well as *calumenin* and *HSP90* as core heat stress response genes in abalone (Barkan et al., 2025). *Formin-binding protein 4-like* (*FNBP4*) likely encodes a cytoskeleton protein that inhibits formin1-mediated actin assembly (Das et al., 2025), which can impact transcription regulation and DNA damage repair (Ulferts et al., 2024; Wollscheid & Ulrich, 2023). Differential *FNBP4* expression has not yet been reported in corals; however, it was recently identified as differentially expressed in response to physiological stress in canines (Leblanc et al., 2021). *Endoplasmin-like isoform X1* (*endoplasmin* or *glucose-regulated protein 94*, *GRP94*; Marzec et al., 2012) likely encodes the endoplasmic reticulum resident paralog of HSP90 (Park et al., 2020) that has not yet been reported in corals following stress exposure; however, a recent study on the marine invertebrate *Batillaria attramentaria* comparing slow versus fast heating to lethal temperature on transcriptome profiles identified *GRP94* as up-regulated under slow-ramp heat stress conditions (Du et al., 2024). Of note is that a protein disulfide isomerase (*PDIA5*) was also up-regulated under fast-ramp heat stress conditions. Lastly, *patched domain-containing protein 3* (*PTCHD3*), which likely encodes a transmembrane protein involved in the developmental Hedgehog signaling pathway (Marigo et al., 1996), was down-regulated in the sea cucumber *Apostichopus japonicus* following exposure to heat stress and low oxygen concurrently and *calreticulin* was up-regulated under heat stress only (Xia et al., 2024). The *A. tersa* recruits in this study exhibited down-regulation of *PTCHD3* following exposure to heat stress conditions, suggesting that 32°C heat stress tanks might have had reduced oxygen relative to 27.5°C ambient tanks (Tromans, 1998). Taken together, these seven genotype-independent heat stress DEGs could be useful molecular markers for monitoring the adaptive heat stress responses in *A. tersa* and other corals targeted for restoration efforts (i.e., quantifiable molecular phenotype to compliment physiological phenotypes), so further testing and function elucidation are warranted.

## Conclusions

SNP genotyping revealed that the utilised broodstock from Martin and Davies reefs exhibited low genetic differentiation (i.e., from same interbreeding population) and showed similar levels of genetic diversity to one another and the four larval offspring groups. Consensus differential expression and GO-slim enrichment analyses revealed that recruits from within- and between-reef offspring groups mounted transcriptional responses at a scale consistent with moderate heat stress, as suggested by the relatively high survival across all recruits (> 65%; Macadam, et al., 2025). These responses involved activation of a largely conserved transcriptional programme consistent with the Type A General Coral Stress Response where both consensus heat stress DEG number and effect size followed D ✕ D > M ✕ D > M ✕ M, which reflects the relative DHW of heat stress experienced among groups. Moreover, seven genotype-independent heat stress responsive candidate genes were consistently identified across all offspring groups, highlighting a conserved molecular core of the coral heat stress response and providing potentially robust targets for thermal stress response monitoring. Taken together, these findings suggest that the benefits of thermal history provenancing-informed selective breeding may be limited in low-differentiation systems and that targeted pre-screening of broodstock may help capture functionally relevant genetic variation relevant to restoration applications.

## Author Contributions

KMQ and MvO obtained the funding and conceptualized the study. KMQ, AM, and CAM designed and ran the experiment. AM, GAM, and RCE produced the genetic data and RCE conducted all analyses. RCE wrote the manuscript with input from CAM, TD, PWL, PB, MvO, and AL.

## Acknowledgements

The authors acknowledge the Traditional Owners of the Great Barrier Reef, particularly the Wulgurukaba and Ngurruumungu people, as custodians of the Sea Country where this work was conducted, and recognise their continuing connection to Country. We thank them for enabling coral collection and breeding within their Sea Country. We pay our respects to their elders past and present and extend that respect to all Aboriginal and Torres Strait Islanders. The authors would also like to acknowledge the Holland Elder (AIMS) for her feedback during analyses, AIMS Marine Operations team for their contributions to successful fieldwork, National Sea Simulator Staff for their help during coral spawning, all AIMS volunteers for their assistance during heat stress experiment, and all AIMS PC2 Lab members for their support during lab work.

## Ethics Statement

Ethics approval is not required for scleractinian coral research in Australia.

## Data Availability Statement

Raw RNAseq and SNP reads are openly available at European Nucleotide Archive (ENA) under project PRJEB103972. This includes RNAseq reads (ERS30577508 – ERS30577547) and SNP reads (ERS30577548 – ERS30577691) included in analyses as well as additional RNAseq reads (ERS30577692 – ERS30577703) and SNP reads (ERS30577704 – ERS30577971) not included in analyses. SNP and RNAseq count matrices, custom R scripts used for SNP and RNAseq analyses, resultant Supplementary Files S1 – S25, and UniProt *BLASTP* result files (*n* = 44) are openly available in AIMS Data Repository at https://doi.org/10.25845/1q96-ja65. *[Please note that data will be made publicly available upon acceptance for publication.]*

