## Supplementary Materials for "Conserved transcriptomic heat stress response signatures in coral recruits selectively bred from thermally distinct broodstock in a low-differentiation system"

### Materials and Methods

#### *Total RNA extraction*

Total RNA was extracted from individual surviving singleton recruits from three Martin ✕ Martin families, three Davies ✕ Davies families, one Davies ✕ Martin family, and two Martin ✕ Davies families using the RNAqueous™- Micro Total RNA Isolation Kit (Invitrogen) kit as follows: (1) RNAlater™ crystal precipitate removed (if present), (2) 6 – 10 Lysis Matrix C beads (MPBio) added to 2 mL screw-top tube containing sample, (3) 100 µL lysis buffer added, (4) homogenized using FastPrep-24 BeadBeater (MPBio) at 5.5 m/s for 30 seconds twice (5 minute pause in between), (5) spun down (pulse), (6) 50 µL absolute ethanol added to lysate, vortexed (10 seconds on maximum), and spun down (pulse), (7) total RNA bound to column membrane (20,100 x g for 1 minute), (8) column membrane washed three times using 180 µL each time (16,500 x g for 30 seconds), (9) column membrane dried (maximum speed for 1 minute), (10) total RNA eluted twice with 7.5 µL elution buffer (70°C), 1 minute at room temperature, and 16,500 x g for 1 minute for each elution (≈ 13 µL total elution). All centrifugations were done in "soft" mode. Extracted total RNA underwent QA/QC using a Bioanalyzer 2100 (Agilent) with RNA 6000 Pico Kit chips (Agilent) by diluting a 1 µL aliquot of each sample by 1:10 – 1:50 in nuclease-free water, denaturing (70°C for 2 minutes), and loading 1 µL of denatured dilution onto individual RNA Pico chips (*n* = 11 samples per chip) along with manufacturer-supplied RNA-specific dye (containing internal standard) and RNA ladder to attain data regarding sample quantity (ng/µL) and integrity (RNA Integrity Number, RIN).

#### *RNA sequencing*

The number of polymerase chain reaction (PCR) cycles applied to each sample during library preparation was adjusted (13 – 15) according to RNA sample concentration gradient. The NEBNext Ultra II library preparation kit was selected because it permits the lowest amount of input RNA (5 ng total) out of the library preparation kits available for non-single-cell sequencing.

*SNP analysis pipeline*

The following sequential filtrations steps were applied to the post-pipeline SNP data delivered by *DArT* for broodstock and larvae (*n* = 144 total samples; Supplemental Table 1) using the package *dartR* version 2 (Mijangos et al., 2022) in *R*: (1) loci with read depth < 5 or > 200 were excluded to reduce the genotype calling error rate; (2) loci with average repeatability < 0.95 were removed to ensure each dataset consisted of loci with high technical repeatability of the sequencing reaction; (3) loci with call rates < 0.80 were removed to reduce missing data; (4) individuals with call rates < 0.75 were removed to further reduce missing data; (5) monomorphic loci were removed due to being invariable; (6) loci with minor allele frequencies < 0.02 were removed to reduce noise from rare variants; and (7) loci out of Hardy-Weinberg equilibrium were removed to avoid confoundment (see Supplemental Files for *R* script).

### Results

*SNPs: larval and adult samples PCoA*

Six of the nine M × M larval families (Supplemental Table 1) formed three distinct and non-overlapping clusters, reflecting the fact that parents are not shared among clusters but both parents are shared within clusters, while one M × M family (M14 × M17) clustered with the M × D family that shared a dam (M14 × D8). The remaining two M × M families (M14 × M9 and M14 × M3) did not cluster together despite sharing a dam; however, this appears to be driven by the large degree of genetic differentiation amongst the M3, M9, and M14 broodstock since each offspring group sits halfway between the parental colonies (Figure 3B). Four of the seven D × D families clustered together, consistent with their half-sibling relationship, while two D × D families clustered together despite no shared parent (D8 × D15 and D12 × D9), and one D × D family (D6 × D15) clustered with a D × M family (D16 × M8) and most D broodstock. One of four M × D families was genetically distinct (M13 × D13), while the other three clustered with the M × M or D × M families that shared one parent, consistent with their half-sibling relationship. Finally, three of four D × M families clustered with M × M, D × D, or M × D families that shared one parent, consistent with their half-sibling relationship, while one D × M family (D16 × M8) clustered with two D × D families despite not sharing a parent (D6 × D15 and D13 × D7).

**Supplementary Figures**

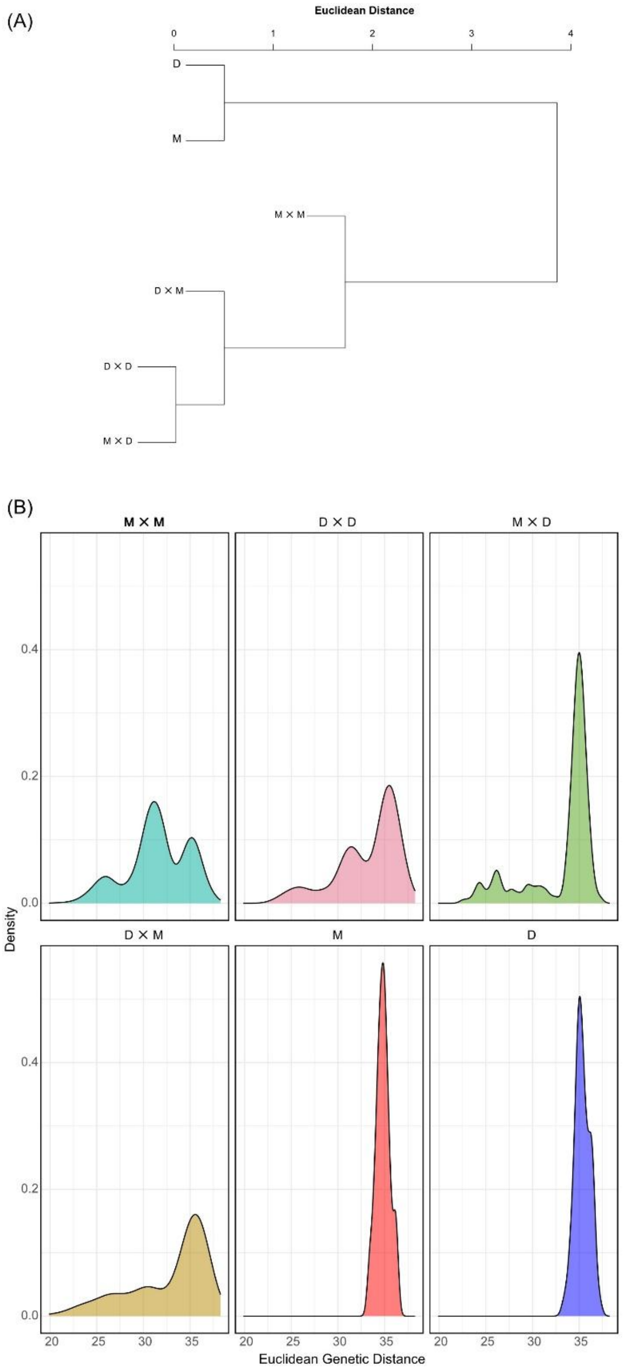


**Supplementary Figure S1.** Dendrogram showing Euclidian genetic distances (i.e., relatedness) amongst Martin (M) and Davies (D) broodstock as well as larvae from within-reef (Martin x Martin, M × M and Davies x Davies, D × D) and between-reef (Martin × Martin, M × M and Davies × Davies, D × D) and between-reef (Martin × Davies, M × D and Davies × Martin, D × M) offspring groups using a filtered SNP dataset (2,337 loci; panel A). Faceted density plot showing Euclidian genetic distances for M and D broodstock as well as each recruit offspring group (B). Labels list dam colony followed by sire colony (separated by '×').

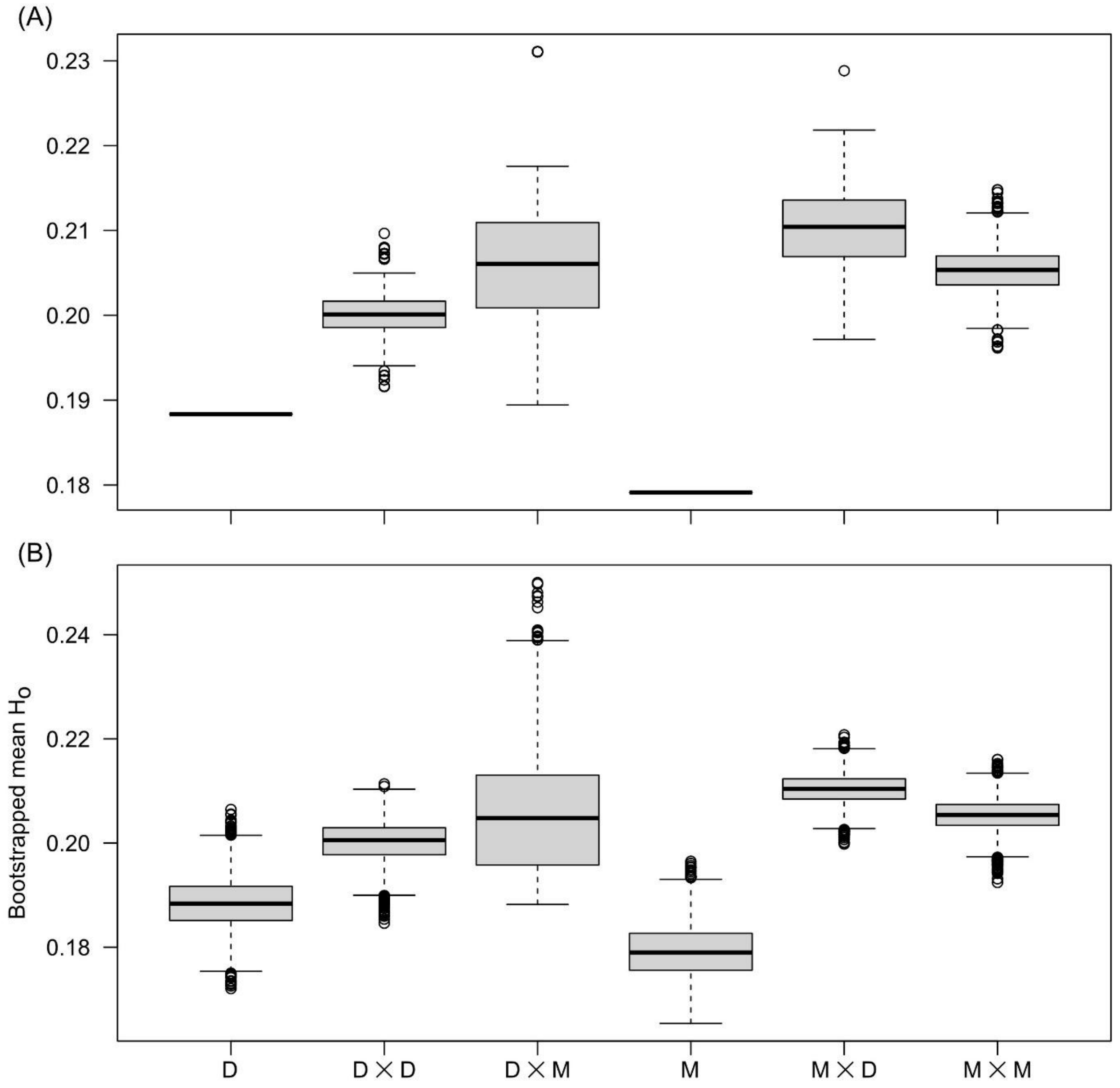


**Supplementary Figure S2.** Boxplots showing variation in observed heterozygosity ($H_o$) following robustness assessment using 5,000 bootstrap replicates at the family-level (A) and individual-level (B). $H_o$ was calculated for broodstock from Martin (M) and Davies (D) reefs as well as larvae from within-reef (Martin × Martin, M × M and Davies × Davies, D × D) and between-reef (Martin × Davies, M × D and Davies × Martin, D × M) offspring groups, using a filtered SNP dataset (2,337 loci). The relatively low variance in family- and individual-level bootstrap distributions indicates that $H_o$ values (Figure 3) are robust to differences in both the number of families and larvae contributing to each offspring group. Labels list dam colony followed by sire colony (separated by '×').

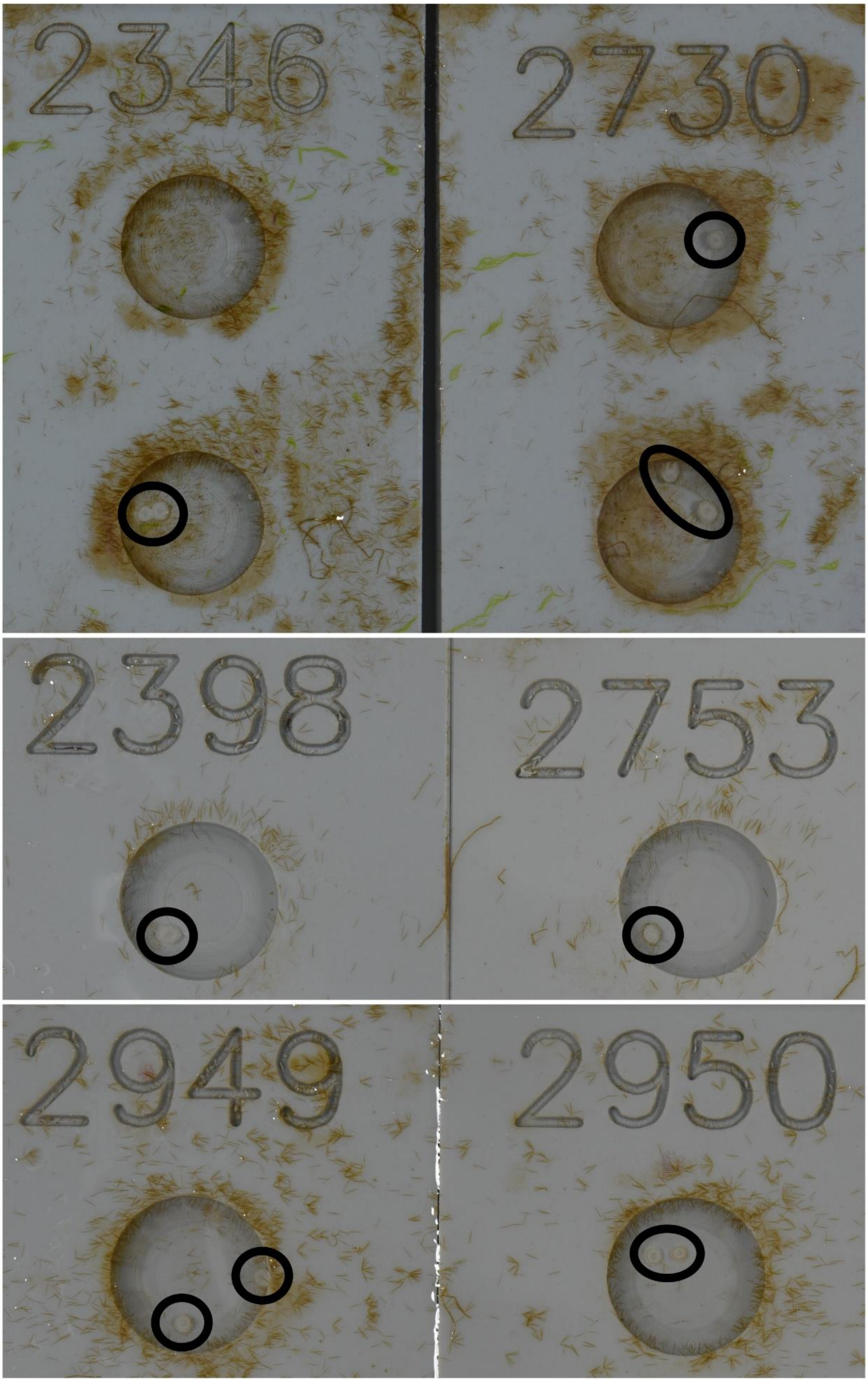


**Supplementary Figure S3.** Images of *Acropora hyacinthus* “neat” recruits (in black circles) from the last time-pointed surveyed (approx. 48 hours before RNA sampling) showing pale colouration (i.e., no symbiont uptake from Wood Reef sediment; see Figure 1C).

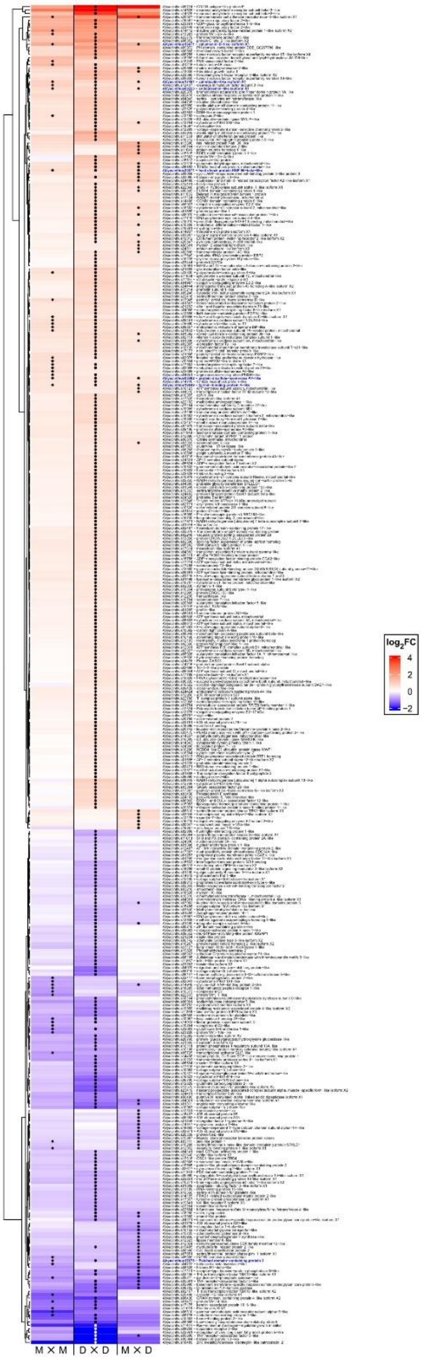


**Supplementary Figure S4.** Heat map of all differentially expressed genes in *Acropora tersa* recruits under heat stress (32°C) relative to ambient (27.5°C) conditions. Rows represent significant differentially expressed genes (DEGs) with $\log_2$ fold-change $\geq 1$ or $\leq -1$ and functional annotations in Martin × Martin (M × M), Davies × Davies (D × D), and Martin × Davies (M × D) recruit offspring groups ($n$ = 12, 14, and 14 averaged individuals respectively). Asterisks denote DEGs that were identified as significant by *edgeR*, *limma voom*, and *DESeq2* (i.e., consensus heat stress DEGs; Benjamini-Hochberg adjusted $p$ value < 0.05). Bolded blue text indicates DEGs that exhibited significantly altered expression in all three recruit offspring groups (i.e., genotype-independent consensus heat stress DEGs). Shades of red and blue represent the magnitude of up- and down-regulation, respectively. Offspring group labels list dam colony followed by sire colony (separated by ‘×’).

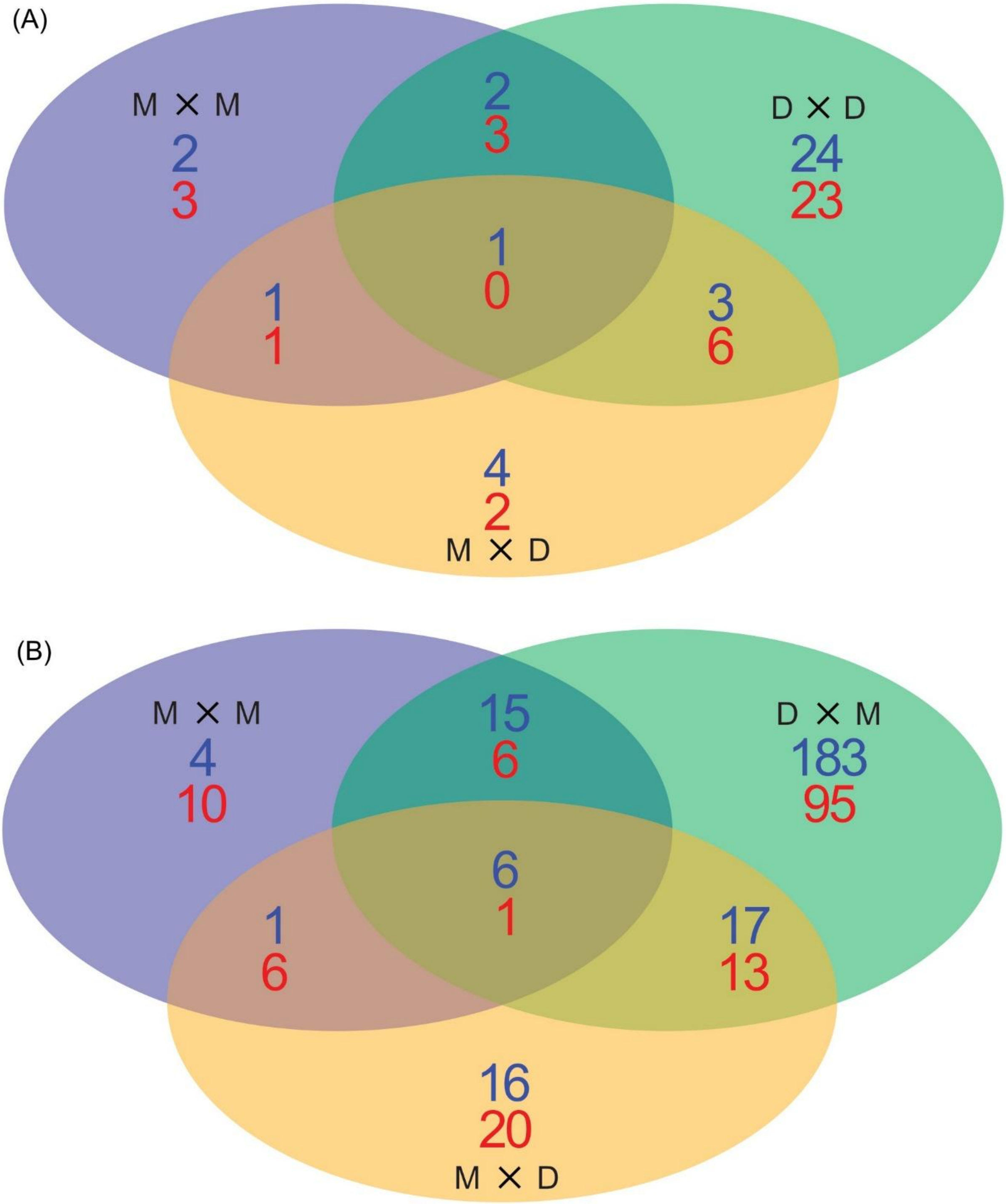


**Supplementary Figure S5.** Venn diagrams showing overlap of significant differentially expressed genes (DEGs) identified by *edgeR*, *limma voom*, and *DESeq2* (i.e., consensus heat stress DEGs; Benjamini-Hochberg adjusted *p* values < 0.05) for recruits from Martin × Martin (M × M), Davies × Davies (D × D), and Martin × Davies (M × D) offspring groups after filtering for: (A) $\log_2$ fold-change > 1 and functional annotations ($n$ = 75 DEGs) and (B) functional annotations only ($n$ = 393 DEGs). Note that panels A and B relate to the DEGs presented in Figure 5 and Supplemental Figure 3 heatmaps, respectively. Top blue and bottom red numbers refer to up- and down-regulated DEGs. Labels list the dam colony followed by the sire colony (separated by '×').

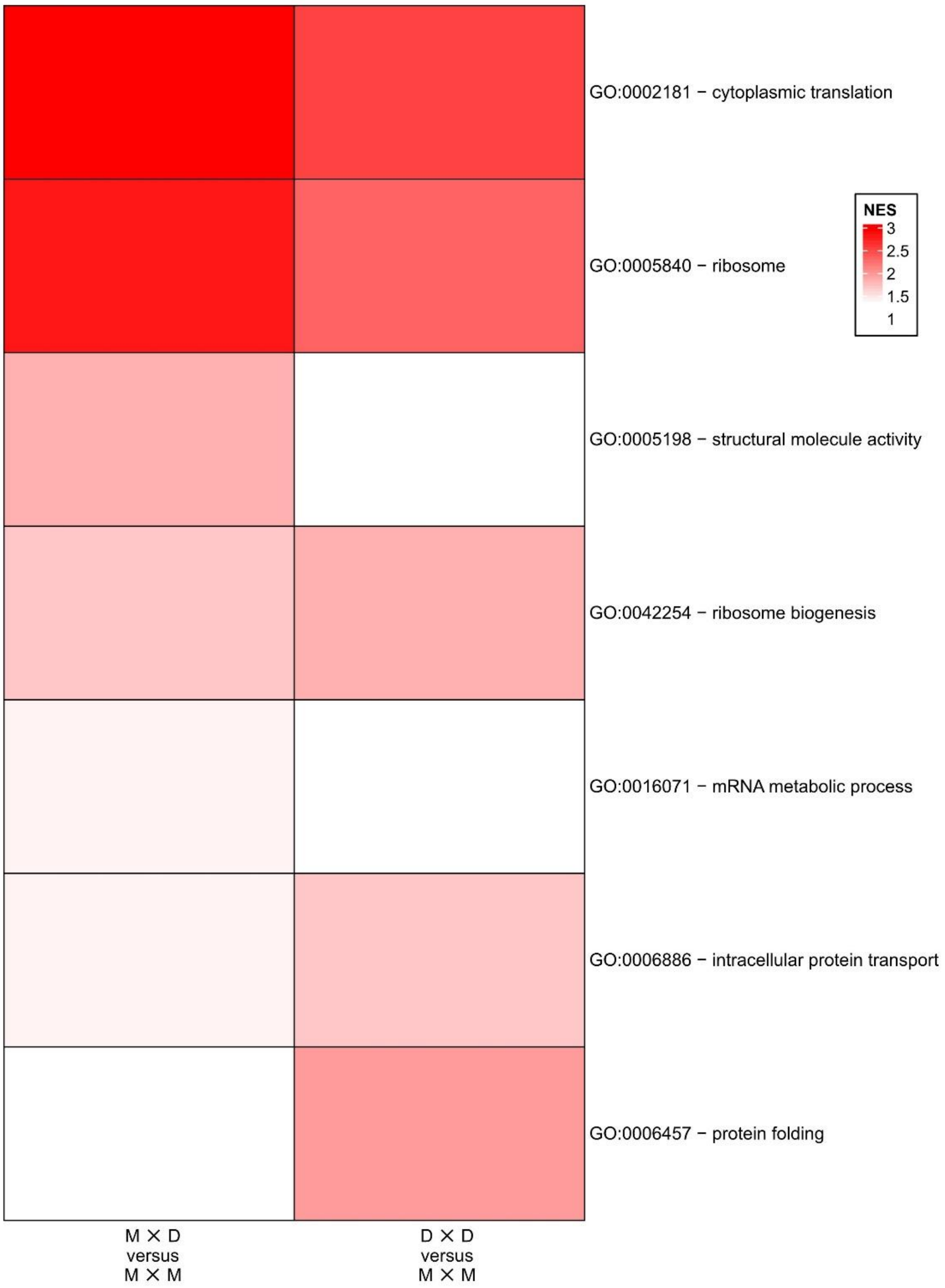


**Supplementary Figure S6.** Heat map of gene ontology (GO)-slim generic subset term enrichment of *Acropora tersa* recruits that exhibited significant enrichment in at least one frontloading comparison among offspring groups (e.g., Martin ╳ Davies [M x D] ambient – Martin ╳ Martin [M x M] ambient). Note that Martin ╳ Davies versus Davies ╳ Davies (D x D) is not included because this comparison yielded no significantly enriched GO slim generic terms (see Table 4). Rows represent significantly enriched GO-slim terms identified in at least one frontloading comparison using gene set enrichment analysis (GSEA) conducted on the regularized log transformed counts per million (CPM) filtered dataset ($n$ = 12,531) for M ╳ M, D ╳ D, and M ╳ D recruits ($n$ = 12, 14, and 14 averaged individuals respectively). Colours represent normalized enrichment score (NES) values (≥ 1.3 or ≤ -1.3). Offspring group labels list dam colony followed by sire colony (separated by '╳').

**Supplementary Tables**

**Supplementary Table S1.** Summary of the number of samples (*n*) sent to service providers for reduce representation whole genome single nucleotide polymorphism sequencing (SNP) and total RNA sequencing (RNAseq) for broodstock from Martin (M) and Davies (D) reefs as well as larvae from within-reef (Martin × Martin, M × M and Davies × Davies, D × D) and between-reef (Martin × Davies, M × D and Davies × Martin, D × M) offspring groups. Bolded families were removed from SNP analysis due having no larvae from heat stress treatment represented. Italicized families were included in recruit heat stress experiment but not included in RNAseq analysis due to insufficient RNA quality or representation. Numbers refer to the unique parental colony ID.

| Family | Stage | SNP (*n*) | RNAseq (*n*) |
|---|---|---|---|
| M | Adult | 9 | 0 |
| D | Adult | 11 | 0 |
| M11 × M14 | Larvae | 6 | 0 |
| M11 × M9 | Larvae | 4 | 0 |
| M13 × M9 | Larvae | 5 | 0 |
| M14 × M11 | Larvae | 5 | 0 |
| M14 × M17 | Larvae | 4 | 0 |
| M14 × M3 | Larvae/Recruit | 5 | 6 |
| M14 × M9 | Larvae | 5 | 0 |
| M9 × M11 | Larvae | 5 | 0 |
| M9 × M13 | Larvae | 5 | 0 |
| M9 × M14 | Recruit | 0 | 6 |
| D12 × D9 | Larvae | 5 | 0 |
| D13 × D14** | Larvae | 4 | 0 |
| D13 × D7 | Larvae | 5 | 0 |
| D6 × D15 | Larvae | 5 | 0 |
| D8 × D12 | Larvae | 6 | 0 |
| D8 × D14 | Larvae/Recruit | 5 | 8 |
| D8 × D15 | Larvae | 5 | 0 |
| D16 × D17 | Recruit | 0 | 6 |
| *D15 × D16* | *Recruit* | *0* | *0* |
| *D13 × D8* | *Recruit* | *0* | *0* |
| *D8 × D13* | *Recruit* | *0* | *0* |
| D13 × M17 | Larvae | 5 | 0 |
| D16 × M8 | Larvae | 5 | 0 |
| D9 × M11 | Larvae | 5 | 0 |
| D10 × M17 | Larvae | 5 | 0 |
| *D6 × M2* | *Recruit* | *0* | *0* |
| *D12 × M15* | *Recruit* | *0* | *0* |
| M11 × D15* | Larvae | 4 | 0 |

| | | | |
|---|---|---|---|
| M11 × D9 | Larvae | 6 | 0 |
| M13 × D13* | Larvae | 5 | 0 |
| M14 × D8 | Larvae | 5 | 0 |
| M17 × D14 | Larvae | 5 | 0 |
| M15 × D12 | Recruit | 0 | 8 |
| M2 × D12 | Recruit | 0 | 6 |
| *M17 × D7* | *Recruit* | *0* | *0* |

*Denotes the number of samples removed during SNP filtering

**Supplementary Table S2.** Summary of RNA quality and quantity sent to service provider for each singleton *Acropoa tersa* recruit from within-reef (Martin × Martin, M × M and Davies × Davies, D × D) and between-reef (Martin × Davies, M × D and Davies × Martin, D × M) offspring groups generated from Martin (M) and/or Davies (D) Reef broodstock. Bolded individual was removed from analysis due to library size < 2.5M.

| Treatment | Cross | RIN | Conc. (ng/µl) | Volume Sent (µl) | Total Sent (ng) |
|---|---|---|---|---|---|
| Ambient | M14 × M3 | 7.2 | 2.78 | 13 (all) | 36.14 |
| Ambient | M14 × M3 | 7.5 | 14.11 | 6.00 | 84.68 |
| Ambient | M 14× M3 | 7.1 | 4.45 | 13 (all) | 57.79 |
| Ambient | M9 × M14 | 7.3 | 5.75 | 6.00 | 34.50 |
| Ambient | M9 × M14 | 7.2 | 17.28 | 6.00 | 103.69 |
| Ambient | M9 × M14 | 6.1 | 6.42 | 6.00 | 38.55 |
| Ambient | D8 × D14 | 6.5 | 9.16 | 6.00 | 54.97 |
| Ambient | D8 × D14 | 6.3 | 8.88 | 6.00 | 53.27 |
| Ambient | D8 × D14 | N/A | 2.40 | 13 (all) | 31.14 |
| Ambient | D8 × D14 | 7.7 | 2.85 | 13 (all) | 37.10 |
| Ambient | D16 × D17 | 7.1 | 4.27 | 13 (all) | 55.53 |
| Ambient | D16 × D17 | 7.2 | 4.83 | 13 (all) | 62.77 |
| Ambient | D16 × D17 | 7.5 | 5.80 | 6.00 | 34.77 |
| Ambient | M15 × D12 | 5.4 | 1.66 | 13 (all) | 21.62 |
| Ambient | M15 × D12 | 7.4 | 2.59 | 13 (all) | 33.70 |
| Ambient | M15 × D12 | 6.8 | 0.99 | 13 (all) | 12.84 |
| Ambient | M15 × D12 | 7.7 | 2.25 | 13 (all) | 29.26 |
| Ambient | M2 × D12 | 7 | 11.36 | 6.00 | 68.16 |
| Ambient | M2 × D12 | 7 | 3.20 | 13 (all) | 41.62 |
| Ambient | M2 × D12 | 6.9 | 13.85 | 6.00 | 83.09 |
| Hot | M14 × M3 | 7 | 11.30 | 6.00 | 67.77 |
| Hot | M14 × M3 | 6.9 | 5.79 | 6.00 | 34.73 |
| Hot | M14 × M3 | 6.8 | 3.75 | 13 (all) | 48.69 |
| Hot | M9 × M14 | N/A | 6.09 | 6.00 | 36.56 |
| Hot | M9 × M14 | 7 | 7.21 | 6.00 | 43.26 |
| Hot | M9 × M14 | 6.6 | 8.13 | 6.00 | 48.75 |
| Hot | D8 × D14 | 7.3 | 3.23 | 13 (all) | 41.95 |
| **Hot** | **D8 × D14** | **7.2** | **3.47** | **13 (all)** | **45.11** |
| Hot | D8 × D14 | 6.5 | 13.14 | 6.00 | 78.82 |
| Hot | D8 × D14 | 6.5 | 5.42 | 6.00 | 32.51 |
| Hot | D16 × D17 | 6.7 | 3.79 | 13 (all) | 49.24 |
| Hot | D16 × D17 | 6.7 | 4.46 | 13 (all) | 57.97 |
| Hot | D16 × D17 | 4.9 | 1.70 | 13 (all) | 22.11 |
| Hot | M15 × D12 | 5.1 | 1.97 | 13 (all) | 25.59 |
| Hot | M15 × D12 | N/A | 9.99 | 6.00 | 59.92 |
| Hot | M15 × D12 | N/A | 13.47 | 6.00 | 80.84 |

| | | | | | |
|---|---|---|---|---|---|
| Hot | M15 × D12 | 6.5 | 2.72 | 13 (all) | 35.39 |
| Hot | M2 × D12 | N/A | 8.93 | 6.00 | 53.58 |
| Hot | M2 × D12 | 6.7 | 17.23 | 6.00 | 103.36 |
| Hot | M2 × D12 | 5.5 | 4.31 | 13 (all) | 56.07 |

**Supplementary Table S3.** Original and bootstrap-derived mean observed heterozygosity ($H_o$) values for *Acropora tersa* broodstock and offspring groups. Original mean $H_o$ (± SD) and corresponding family- and individual-level bootstrap estimates (mean and 95% confidence intervals; 5,000 replicates) were calculated for broodstock from Martin (M) and Davies (D) reefs, as well as larvae from within-reef (M ╳ M, D ╳ D) and between-reef (M ╳ D, D ╳ M) offspring groups, using the filtered SNP dataset (2,337 loci). Original mean $H_o$ values were used in one-way analysis of variance (ANOVA) with Tukey's HSD post-hoc (Figure 3). Sample sizes (*n*) for each group are shown in parentheses. Labels list dam colony followed by sire colony (separated by '╳').

| | Original (ANOVA) | | Family-level Bootstrap | | | Individual-level Bootstrap | | |
|---|---|---|---|---|---|---|---|---|
| | mean_Ho | SD_Ho | mean_Ho | CI_low | CI_high | mean_Ho | CI_low | CI_high |
| M ╳ M (*n* = 44) | 0.205 | 0.021 | 0.205 | 0.200 | 0.210 | 0.205 | 0.199 | 0.211 |
| M ╳ D (*n* = 23) | 0.210 | 0.014 | 0.210 | 0.202 | 0.218 | 0.210 | 0.205 | 0.216 |
| D ╳ M (*n* = 20) | 0.206 | 0.049 | 0.206 | 0.194 | 0.218 | 0.205 | 0.192 | 0.229 |
| D ╳ D (*n* = 33) | 0.200 | 0.022 | 0.200 | 0.195 | 0.205 | 0.200 | 0.192 | 0.207 |
| D (*n* = 11) | 0.188 | 0.017 | 0.188 | 0.188 | 0.188 | 0.188 | 0.179 | 0.198 |
| M (*n* = 9) | 0.179 | 0.016 | 0.179 | 0.179 | 0.179 | 0.179 | 0.170 | 0.190 |

**Supplementary Table S4**. Differentially expressed genes with functional annotations and $log_2$ fold-change (log2FC) > 1 (i.e., fold-change > 2) that were identified by *edgeR*, *limma voom*, and *DESeq2* for recruits from within-reef (Martin ╳ Martin, M ╳ M and Davies ╳ Davies, D ╳ D) and between-reef (Martin ╳ Davies, M ╳ D) offspring groups subjected to elevated temperature conditions. Labels list dam colony followed by sire colony (separated by '╳').

| | | | | *edgeR* | | *limma voom* | | *DESeq2* | |
|---|---|---|---|---|---|---|---|---|---|
| **Cross** | **Direction** | **Annotation** | **geneID** | **log2FC** | **adj.P.Val** | **log2FC** | **adj.P.Val** | **log2FC** | **adj.P.Val** |
| M ╳ M | UP | FAS-associated factor 2-like | Ahyacinthus19245 | 1.3 | 9.60E-03 | 1.3 | 1.50E-03 | 1.3 | 1.40E-04 |
| | | transmembrane cell adhesion receptor mua-3-like isoform X1 | Ahyacinthus20027 | 1.8 | 9.60E-03 | 1.8 | 1.40E-02 | 1.9 | 4.40E-02 |
| | | tubulin beta-4B chain | Ahyacinthus01218 | 1.4 | 2.10E-03 | 1.4 | 6.70E-04 | 1.5 | 6.30E-05 |
| | | protein Wnt-2b-A-like | Ahyacinthus13383 | 1.4 | 2.20E-02 | 1.5 | 8.00E-03 | 1.4 | 1.50E-03 |
| | | inactive pancreatic lipase-related protein 1-like isoform X2 | Ahyacinthus14412 | 1.9 | 2.00E-02 | 1.9 | 4.80E-02 | 2.1 | 4.30E-02 |
| | | calumenin-B-like isoform X1 | Ahyacinthus19471 | 1.7 | 1.80E-02 | 1.7 | 3.40E-04 | 1.7 | 1.10E-03 |
| | DOWN | gamma-aminobutyric acid receptor subunit alpha-5-like | Ahyacinthus08510 | -1.8 | 1.20E-02 | -1.8 | 1.80E-02 | -1.8 | 4.10E-02 |
| | | protein Wnt-4-like | Ahyacinthus01472 | -1.2 | 1.70E-02 | -1.2 | 1.40E-02 | -1.3 | 1.70E-03 |
| | | Protein very KIND | Ahyacinthus11036 | -1.2 | 4.80E-02 | -1.2 | 2.90E-02 | -1.2 | 4.70E-02 |
| | | protein Wnt-1-like | Ahyacinthus22633 | -1 | 9.60E-03 | -1 | 1.90E-03 | -1 | 4.80E-04 |
| | | collectin-12-like isoform X1 | Ahyacinthus20408 | -1.5 | 4.90E-02 | -1.5 | 1.90E-02 | -1.5 | 5.80E-03 |
| | | endothelin-converting enzyme 1-like | Ahyacinthus06776 | -1.5 | 1.70E-02 | -1.6 | 2.00E-03 | -1.5 | 4.30E-02 |
| | | pyridoxine-5'-phosphate oxidase-like | Ahyacinthus05211 | -1 | 2.60E-02 | -1 | 9.80E-03 | -1 | 1.10E-02 |
| D ╳ D | UP | matrix metalloproteinase-2-like | Ahyacinthus20888 | 1.2 | 1.70E-02 | 1.1 | 1.80E-02 | 1.2 | 9.80E-04 |
| | | tubulin beta-4B chain | Ahyacinthus01218 | 1.1 | 1.70E-02 | 0.9 | 2.30E-02 | 1.1 | 3.40E-03 |
| | | coactosin-like protein | Ahyacinthus15812 | 1.1 | 2.90E-02 | 1 | 2.00E-02 | 1.1 | 1.10E-02 |
| | | glia maturation factor beta-like | Ahyacinthus21255 | 1 | 1.50E-02 | 0.9 | 3.10E-02 | 1.1 | 2.10E-03 |
| | | UDP-glucuronosyltransferase 1-5-like | Ahyacinthus23261 | 2.2 | 2.20E-02 | 2.3 | 2.00E-03 | 2.2 | 9.80E-04 |
| | | serine--pyruvate aminotransferase-like | Ahyacinthus04849 | 1.2 | 1.50E-02 | 1.1 | 6.00E-03 | 1.2 | 1.70E-03 |
| | | alkaline phosphatase-like | Ahyacinthus04735 | 1.1 | 3.90E-03 | 1.1 | 1.70E-03 | 1.1 | 4.40E-04 |
| | | glutamate dehydrogenase, mitochondrial-like | Ahyacinthus12318 | 1 | 2.00E-02 | 1 | 5.40E-03 | 1 | 4.00E-03 |
| | | NADH dehydrogenase [ubiquinone] 1 alpha subcomplex subunit 13-like | Ahyacinthus08849 | 1 | 2.40E-03 | 1 | 4.70E-04 | 1 | 3.70E-04 |
| | | DBH-like monooxygenase protein 1 | Ahyacinthus03883 | 1 | 3.70E-03 | 1.1 | 7.20E-04 | 1 | 2.10E-04 |
| | | neuronal acetylcholine receptor subunit beta-3-like | Ahyacinthus14528 | 4.7 | 1.00E-03 | 5.4 | 2.30E-02 | 4.2 | 8.30E-03 |

| | | | | | | | | |
|---|---|---|---|---|---|---|---|---|
| | neuronal acetylcholine receptor subunit beta-3-like | Ahyacinthus14529 | 3.8 | 2.40E-03 | 3.8 | 6.30E-03 | 3.3 | 8.30E-03 |
| | protein C-ets-2-like isoform X2 | Ahyacinthus08352 | 2.4 | 2.30E-02 | 2.2 | 4.40E-02 | 2.4 | 1.50E-03 |
| | PH domain-containing protein DDB_G0267786-like | Ahyacinthus09972 | 2.1 | 5.10E-03 | 1.9 | 6.00E-03 | 2.2 | 6.60E-06 |
| | transmembrane protein 205-like | Ahyacinthus22779 | 2 | 2.10E-02 | 1.8 | 4.10E-02 | 1.9 | 2.20E-02 |
| | protein Wnt-2b-A-like | Ahyacinthus13383 | 1.9 | 1.00E-03 | 1.9 | 1.00E-03 | 1.9 | 5.10E-08 |
| | sterile alpha motif domain-containing protein 11-like | Ahyacinthus03949 | 1.8 | 1.40E-02 | 1.8 | 2.80E-02 | 1.6 | 1.50E-02 |
| | fibroblast growth factor 1-like | Ahyacinthus06868 | 1.6 | 2.90E-02 | 1.5 | 3.60E-02 | 1.6 | 2.00E-02 |
| | Fibroblast growth factor 1 | Ahyacinthus13035 | 1.4 | 3.70E-03 | 1.4 | 9.80E-04 | 1.4 | 1.60E-04 |
| | protein Wnt-2b-A-like | Ahyacinthus13382 | 1.3 | 4.90E-02 | 1.2 | 3.70E-02 | 1.2 | 1.60E-02 |
| | fibroblast growth factor receptor 3-like isoform X2 | Ahyacinthus23595 | 1.3 | 2.80E-02 | 1.4 | 4.30E-02 | 1.7 | 9.00E-03 |
| | sterile alpha motif domain-containing protein 11-like | Ahyacinthus03950 | 1.1 | 3.80E-02 | 1.1 | 3.40E-02 | 1.1 | 2.00E-02 |
| | CD209 antigen-like protein E | Ahyacinthus05724 | 5.9 | 1.20E-04 | 5 | 7.60E-03 | 5.9 | 1.00E-07 |
| | interferon regulatory factor 2-like | Ahyacinthus16587 | 2.4 | 1.20E-02 | 2.8 | 4.70E-02 | 2.5 | 1.20E-02 |
| | interferon regulatory factor 2-like | Ahyacinthus22448 | 2.1 | 1.20E-02 | 2.1 | 6.60E-03 | 2.2 | 5.00E-04 |
| | tumor necrosis factor receptor superfamily member 1B-like isoform X4 | Ahyacinthus08258 | 1.6 | 2.60E-02 | 1.7 | 7.40E-03 | 1.6 | 4.00E-03 |
| | stimulator of interferon genes protein-like | Ahyacinthus11339 | 1.4 | 3.50E-02 | 1.5 | 1.50E-02 | 1.4 | 2.00E-02 |
| | tumor necrosis factor receptor superfamily member 19-like | Ahyacinthus09850 | 1.3 | 7.80E-03 | 1.4 | 1.40E-03 | 1.3 | 4.40E-04 |
| | baculoviral IAP repeat-containing protein 5-like | Ahyacinthus18512 | 1.3 | 2.80E-02 | 1.5 | 1.50E-02 | 1.3 | 4.80E-02 |
| | cathepsin Z-like | Ahyacinthus12192 | 1 | 1.20E-04 | 0.9 | 2.30E-04 | 1 | 2.00E-08 |
| | calumenin-B-like isoform X1 | Ahyacinthus19471 | 1.6 | 7.40E-03 | 1.7 | 2.00E-04 | 1.6 | 4.80E-04 |
| | 10 kDa heat shock protein, mitochondrial-like | Ahyacinthus06892 | 1 | 1.00E-03 | 1 | 2.80E-04 | 1 | 1.20E-05 |
| | bifunctional arginine demethylase and lysyl-hydroxylase JMJD6-like | Ahyacinthus18259 | 1.2 | 3.90E-02 | 1.4 | 3.90E-02 | 1.2 | 9.30E-03 |
| | polyadenylate-binding protein 4-like | Ahyacinthus22329 | 1.1 | 6.80E-03 | 1 | 4.60E-03 | 1.1 | 3.50E-04 |
| DOWN | dopamine receptor 3-like | Ahyacinthus08259 | -2.2 | 6.30E-03 | -2.3 | 4.30E-02 | -2.1 | 6.40E-03 |
| | Ras-related and estrogen-regulated growth inhibitor | Ahyacinthus11404 | -2 | 1.20E-02 | -1.9 | 3.50E-02 | -1.9 | 2.10E-02 |
| | TNF receptor-associated factor 5-like | Ahyacinthus05600 | -1.9 | 3.70E-03 | -2.1 | 3.90E-02 | -2.3 | 7.10E-06 |
| | Low-density lipoprotein receptor-related protein 6 | Ahyacinthus09989 | -1.7 | 3.50E-03 | -1.5 | 2.00E-02 | -1.8 | 1.20E-05 |

| | | | | | | | |
|---|---|---|---|---|---|---|---|
| GRAM domain-containing protein 4-like isoform X2 | Ahyacinthus08430 | -1.7 | 2.70E-02 | -1.8 | 2.90E-02 | -1.6 | 4.80E-03 |
| protein Wnt-4-like | Ahyacinthus01472 | -1.4 | 2.90E-03 | -1.2 | 6.00E-03 | -1.4 | 2.70E-05 |
| T-box transcription factor TBX10-like isoform X2 | Ahyacinthus02157 | -1.4 | 1.70E-02 | -1.3 | 4.00E-02 | -1.4 | 1.40E-02 |
| endothelin-converting enzyme 1-like | Ahyacinthus06776 | -1.3 | 1.70E-02 | -1.4 | 5.70E-03 | -1.3 | 2.50E-02 |
| angiotensin-converting enzyme-like | Ahyacinthus01311 | -1.2 | 3.90E-03 | -1.2 | 4.10E-03 | -1.4 | 6.20E-05 |
| endothelin-converting enzyme homolog isoform X1 | Ahyacinthus04021 | -1.1 | 7.40E-03 | -1 | 1.40E-02 | -1.2 | 9.80E-04 |
| TNF receptor-associated factor 5-like | Ahyacinthus05601 | -1 | 2.90E-02 | -0.9 | 4.60E-02 | -1.1 | 5.80E-03 |
| transcription factor Sp9-like | Ahyacinthus24451 | -1 | 2.30E-02 | -1.2 | 9.40E-03 | -1 | 1.40E-02 |
| elongation of very long chain fatty acids protein 5-like | Ahyacinthus22861 | -2.1 | 2.90E-02 | -2.8 | 4.40E-02 | -2.3 | 3.40E-03 |
| heme-binding protein 2-like | Ahyacinthus10383 | -1.5 | 4.20E-02 | -1.3 | 4.20E-02 | -1.5 | 1.40E-02 |
| glutamate carboxypeptidase 2-like | Ahyacinthus12629 | -1.3 | 3.00E-03 | -1.2 | 5.20E-03 | -1.3 | 1.00E-04 |
| cytochrome P450 4F4-like | Ahyacinthus16323 | -1.3 | 2.50E-02 | -1.1 | 3.90E-02 | -1.3 | 7.40E-03 |
| D-arabinono-1,4-lactone oxidase | Ahyacinthus08782 | -1.2 | 2.40E-02 | -1.6 | 7.60E-03 | -1.1 | 4.50E-02 |
| heparan sulfate glucosamine 3-O-sulfotransferase 5-like | Ahyacinthus06147 | -1.2 | 1.20E-02 | -1.4 | 1.50E-03 | -1.1 | 7.40E-03 |
| putative N-acetylated-alpha-linked acidic dipeptidase isoform X1 | Ahyacinthus06496 | -1.1 | 6.60E-03 | -1.1 | 2.60E-03 | -1.1 | 1.40E-03 |
| heparan-alpha-glucosaminide N-acetyltransferase-like | Ahyacinthus10227 | -1.1 | 2.10E-02 | -1 | 1.10E-02 | -1 | 1.20E-02 |
| multidrug resistance-associated protein 4-like isoform X2 | Ahyacinthus19959 | -1.1 | 3.00E-02 | -1 | 3.20E-02 | -1.1 | 2.30E-02 |
| zinc metalloproteinase-disintegrin-like batroxstatin-2 | Ahyacinthus18409 | -2.5 | 7.40E-03 | -2.7 | 2.90E-02 | -2.2 | 3.90E-02 |
| serine-rich adhesin for platelets-like isoform X5 | Ahyacinthus13575 | -1.5 | 2.40E-02 | -1.3 | 3.30E-02 | -1.5 | 4.70E-03 |
| Protein very KIND | Ahyacinthus11036 | -1.4 | 1.50E-02 | -1.3 | 3.70E-02 | -1.5 | 1.40E-03 |
| collagen alpha-1(I) chain-like | Ahyacinthus10063 | -1.4 | 2.00E-03 | -1.4 | 7.20E-04 | -1.4 | 3.30E-05 |
| collagen alpha-1(XII) chain-like | Ahyacinthus06700 | -1.4 | 9.90E-03 | -1.3 | 1.50E-02 | -1.5 | 3.80E-03 |
| collagen alpha-2(I) chain-like | Ahyacinthus06710 | -1.3 | 1.50E-02 | -1.3 | 3.40E-02 | -1.3 | 7.30E-03 |
| basement membrane-specific heparan sulfate proteoglycan core protein-like | Ahyacinthus02990 | -1.3 | 4.00E-03 | -1.2 | 6.00E-03 | -1.3 | 9.00E-04 |
| keratin-associated protein 10-6-like | Ahyacinthus17175 | -1.3 | 4.20E-02 | -1.4 | 4.60E-02 | -1.3 | 3.70E-02 |
| migration and invasion-inhibitory protein-like | Ahyacinthus04070 | -1.3 | 6.30E-03 | -1 | 2.70E-02 | -1.4 | 1.50E-04 |
| serine-rich adhesin for platelets-like | Ahyacinthus08865 | -1.1 | 3.00E-02 | -1.1 | 2.40E-02 | -1.1 | 4.50E-02 |

| | | | | | | | | | |
|---|---|---|---|---|---|---|---|---|---|
| | | kinesin-like protein KIF15 isoform X1 | Ahyacinthus11366 | -1 | 3.30E-02 | -1.1 | 1.10E-02 | -1 | 2.50E-02 |
| | | sodium/glucose cotransporter 5-like | Ahyacinthus08091 | -1 | 3.90E-02 | -0.9 | 4.20E-02 | -1.1 | 5.90E-03 |
| | | balbiani ring protein 3-like | Ahyacinthus00084 | -2.5 | 2.90E-03 | -2.4 | 4.10E-02 | -2.6 | 4.70E-03 |
| | | nascent polypeptide-associated complex subunit alpha, muscle-specific form-like isoform X2 | Ahyacinthus23419 | -1.5 | 3.70E-03 | -1.4 | 5.70E-03 | -1.5 | 6.50E-05 |
| M × D | UP | transmembrane cell adhesion receptor mua-3-like isoform X1 | Ahyacinthus20027 | 3.6 | 2.20E-05 | 3.7 | 4.10E-02 | 3.8 | 1.70E-09 |
| | | protein crumbs homolog 1-like | Ahyacinthus11046 | 1.4 | 9.30E-03 | 1.2 | 1.80E-02 | 1.4 | 1.30E-04 |
| | | matrix metalloproteinase-2-like | Ahyacinthus20888 | 1.2 | 1.90E-02 | 1.3 | 1.80E-02 | 1.2 | 6.40E-04 |
| | | cyclin-dependent kinase 2-like | Ahyacinthus06744 | 1.5 | 9.70E-03 | 1.6 | 4.10E-02 | 1.5 | 1.20E-03 |
| | | fibroblast growth factor 1 | Ahyacinthus13035 | 1.2 | 1.30E-02 | 1 | 3.80E-02 | 1.2 | 1.90E-03 |
| | | doublesex- and mab-3-related transcription factor A2-like isoform X1 | Ahyacinthus24486 | 1.1 | 1.30E-02 | 1 | 4.70E-02 | 1.1 | 1.60E-03 |
| | | ras-related protein Rab-30-like | Ahyacinthus13600 | 1.6 | 7.30E-03 | 1.5 | 4.70E-02 | 1.6 | 5.90E-05 |
| | | glutamate dehydrogenase, mitochondrial-like | Ahyacinthus12318 | 1.1 | 1.80E-02 | 0.9 | 3.30E-02 | 1.1 | 1.90E-03 |
| | | calumenin-B-like isoform X1 | Ahyacinthus19471 | 1.6 | 1.30E-02 | 1.6 | 1.80E-03 | 1.5 | 1.00E-03 |
| | | KDEL motif-containing protein 1-like | Ahyacinthus15615 | 1.3 | 2.70E-02 | 1.8 | 3.60E-02 | 1.2 | 4.10E-02 |
| | DOWN | TNF receptor-associated factor 5-like | Ahyacinthus05600 | -1.3 | 1.50E-02 | -1.4 | 1.80E-02 | -1.4 | 1.60E-02 |
| | | TNF receptor-associated factor 5-like | Ahyacinthus05601 | -1.2 | 8.30E-03 | -1.4 | 2.50E-03 | -1.2 | 1.90E-03 |
| | | basement membrane-specific heparan sulfate proteoglycan core protein-like | Ahyacinthus02990 | -1.2 | 7.30E-03 | -1.1 | 1.80E-02 | -1.4 | 5.40E-04 |
| | | histone H1-delta-like | Ahyacinthus08839 | -1.2 | 3.30E-02 | -1.3 | 4.70E-02 | -1.3 | 1.10E-02 |
| | | pyridoxine-5'-phosphate oxidase-like | Ahyacinthus05211 | -1.4 | 3.30E-04 | -1.5 | 4.70E-04 | -1.4 | 2.30E-07 |
| | | D-arabinono-1,4-lactone oxidase | Ahyacinthus08782 | -1.4 | 9.10E-03 | -1.2 | 4.10E-02 | -1.6 | 6.40E-04 |
| | | mitochondrial glycine transporter-like | Ahyacinthus10152 | -1.1 | 1.40E-02 | -1.1 | 2.00E-02 | -1 | 3.40E-03 |
| | | endothelin-converting enzyme homolog isoform X1 | Ahyacinthus04021 | -1.1 | 9.60E-03 | -0.9 | 1.80E-02 | -1.1 | 1.00E-03 |

**Supplementary Table S5.** UniProt *BLASTP* results for putative adaptive heat stress response candidate genes with multiple identical functional annotations identified in all recruits from both within- and between-reef offspring groups.

| **Reference Annotation** | | **UniProt BLASTP Result** | | | | |
|---|---|---|---|---|---|---|
| geneID | Description | Species | Accession | Description | Identity (%) | E value |
| Ahyacinthus14528* | neuronal acetylcholine receptor subunit beta-3-like | *Acropora cervicornis* | A0AAD9QKX9 | neuronal acetylcholine receptor subunit beta-3 | 77.5% | 0 |
| Ahyacinthus14529* | neuronal acetylcholine receptor subunit beta-3-like | *A. cervicornis* | A0AAD9QKX9 | neuronal acetylcholine receptor subunit beta-3 | 77.8% | 0 |
| Ahyacinthus03949* | sterile alpha motif domain-containing protein 11-like | *A. cervicornis* | A0AAD9PS80 | deformed epidermal autoregulatory factor 1 | 99% | 1.1E-139 |
| Ahyacinthus03950* | sterile alpha motif domain-containing protein 11-like | *A. cervicornis* | A0AAD9PS80 | deformed epidermal autoregulatory factor 1 | 83.9% | 0 |
| Ahyacinthus13382* | protein Wnt-2b-A-like | *A. cervicornis* | A0AAD9VBQ1 | Protein Wnt | 53.9% | 1.7E-129 |
| Ahyacinthus13383* | protein Wnt-2b-A-like | *A. cervicornis* | A0AAD9VBQ1 | Protein Wnt | 98.7% | 0 |
| Ahyacinthus16587* | interferon regulatory factor 2-like | *A. cervicornis* | A0AAD9QF57 | Interferon regulatory factor 4 | 86.7% | 0 |
| Ahyacinthus22448* | interferon regulatory factor 2-like | *A. cervicornis* | A0AAD9PSJ6 | Interferon regulatory factor 1 | 94.3% | 1.1E-159 |
| Ahyacinthus05600* | TNF receptor-associated factor 5-like | *A. cervicornis* | A0AAD9QZ12 | TNF receptor-associated factor 4 | 96.5% | 1.2E-121 |
| Ahyacinthus05601* | TNF receptor-associated factor 5-like | *A. cervicornis* | A0AAD9QZ12 | TNF receptor-associated factor 4 | 97.5% | 6.7E-173 |

*see individual UniProt *BLASTP* result files with same geneID name.

**Supplementary Table S6.** Differentially expressed genes (DEGs) identified by *edgeR*, *limma voom*, and *DESeq2* (i.e., consensus) in *Acropora tersa* recruits under heat stress relative to ambient conditions that were common among within-reef (Martin ╳ Martin, M ╳ M and Davies ╳ Davies, D ╳ D) and between-reef (Martin ╳ Davies, M ╳ D) offspring groups as well as unique to between-reef recruits (i.e., common and M ╳ D unique consensus heat stress DEGs). Labels list dam colony followed by sire colony (separated by '╳').

| Comparison | Direction | Annotation | geneID | log2FC^ | SD |
|---|---|---|---|---|---|
| All offspring groups | UP | calumenin-B-like isoform X1 | Ahyacinthus19471 | 1.6 | 0.1 |
| | | heat shock protein HSP 90-beta-like | Ahyacinthus20571 | 0.8 | 0.1 |
| | | calreticulin-like isoform X1 | Ahyacinthus17491 | 0.8 | 0.2 |
| | | endoplasmin-like isoform X1 | Ahyacinthus08223 | 0.8 | 0.1 |
| | | hypothetical protein pdam_00000357, partial* | Ahyacinthus02976 | 0.7 | 0.1 |
| | | protein disulfide-isomerase A4-like | Ahyacinthus05582 | 0.6 | 0.1 |
| | | formin-binding protein 4-like | Ahyacinthus19849 | 0.5 | 0.1 |
| | DOWN | predicted protein* | Ahyacinthus02626 | -1.3 | 0.3 |
| | | uncharacterized protein LOC114955431* | Ahyacinthus15775 | -1.3 | 0.1 |
| | | Patched domain-containing protein 3 | Ahyacinthus17678 | -0.7 | 0.1 |
| M ╳ M + D ╳ D | UP | calumenin-B-like isoform X1 | Ahyacinthus19471 | 1.7 | 0.05 |
| | | protein Wnt-2b-A-like | Ahyacinthus13383 | 1.7 | 0.3 |
| | | tubulin beta-4B chain | Ahyacinthus01218 | 1.2 | 0.2 |
| | | calreticulin-like isoform X1 | Ahyacinthus17491 | 0.9 | 0.04 |
| | | cleavage stimulation factor subunit 3-like | Ahyacinthus12437 | 0.8 | 0.2 |
| | | endoplasmin-like isoform X1 | Ahyacinthus08223 | 0.8 | 0.1 |
| | | cathepsin Z-like | Ahyacinthus12192 | 0.8 | 0.2 |
| | | heat shock protein HSP 90-beta-like | Ahyacinthus20571 | 0.8 | 0.1 |
| | | peptidyl-prolyl cis-trans isomerase D-like | Ahyacinthus13045 | 0.7 | 0.04 |
| | | polyadenylate-binding protein 4-like | Ahyacinthus22330 | 0.7 | 0.1 |
| | | cytochrome c oxidase subunit NDUFA4-like | Ahyacinthus05461 | 0.7 | 0.1 |
| | | endoplasmic reticulum chaperone BiP-like | Ahyacinthus09857 | 0.7 | 0.01 |
| | | cytochrome b5-like isoform X1 | Ahyacinthus16690 | 0.7 | 0.03 |
| | | hypothetical protein pdam_00000357, partial* | Ahyacinthus02976 | 0.6 | 0.04 |
| | | Cytochrome c oxidase subunit 7A-related protein, mitochondrial | Ahyacinthus21182 | 0.6 | 0.04 |
| | | inosine-uridine preferring nucleoside hydrolase-like | Ahyacinthus02075 | 0.6 | 0.1 |
| | | protein disulfide-isomerase A4-like | Ahyacinthus05582 | 0.6 | 0.05 |
| | | serine/arginine-rich splicing factor 7-like | Ahyacinthus17582 | 0.6 | 0.05 |

| | | | | | |
|---|---|---|---|---|---|
| | | endoplasmic reticulum resident protein 29-like | Ahyacinthus02086 | 0.6 | 0.1 |
| | | formin-binding protein 4-like | Ahyacinthus19849 | 0.5 | 0.03 |
| | | cilia- and flagella-associated protein 44-like isoform X1 | Ahyacinthus01666 | 0.5 | 0.1 |
| | | arginine/serine-rich protein PNISR-like | Ahyacinthus04651 | 0.5 | 0.05 |
| | DOWN | endothelin-converting enzyme 1-like | Ahyacinthus06776 | -1.4 | 0.1 |
| | | uncharacterized protein LOC114955431* | Ahyacinthus15775 | -1.4 | 0.1 |
| | | predicted protein* | Ahyacinthus02626 | -1.4 | 0.3 |
| | | Protein very KIND | Ahyacinthus11036 | -1.3 | 0.1 |
| | | protein Wnt-4-like | Ahyacinthus01472 | -1.3 | 0.1 |
| | | LOW QUALITY PROTEIN: uncharacterized protein LOC114972637* | Ahyacinthus11523 | -0.9 | 0.03 |
| | | glutathione S-transferase 1-like | Ahyacinthus22003 | -0.8 | 0.1 |
| | | linear gramicidin synthase subunit D | Ahyacinthus14043 | -0.7 | 0.1 |
| | | lysyl oxidase homolog 2A-like | Ahyacinthus10987 | -0.7 | 0.1 |
| | | Patched domain-containing protein 3 | Ahyacinthus17678 | -0.7 | 0.02 |
| M × M + M × D | UP | transmembrane cell adhesion receptor mua-3-like isoform X1 | Ahyacinthus20027 | 2.7 | 1.0 |
| | | calumenin-B-like isoform X1 | Ahyacinthus19471 | 1.6 | 0.1 |
| | | heat shock protein HSP 90-beta-like | Ahyacinthus20571 | 0.8 | 0.1 |
| | | calreticulin-like isoform X1 | Ahyacinthus17491 | 0.8 | 0.2 |
| | | endoplasmin-like isoform X1 | Ahyacinthus08223 | 0.7 | 0.1 |
| | | hypothetical protein pdam_00000357, partial* | Ahyacinthus02976 | 0.6 | 0.03 |
| | | protein disulfide-isomerase A4-like | Ahyacinthus05582 | 0.6 | 0.04 |
| | | formin-binding protein 4-like | Ahyacinthus19849 | 0.5 | 0.01 |
| | DOWN | predicted protein* | Ahyacinthus02626 | -1.5 | 0.2 |
| | | uncharacterized protein LOC114955431* | Ahyacinthus15775 | -1.3 | 0.1 |
| | | pyridoxine-5'-phosphate oxidase-like | Ahyacinthus05211 | -1.2 | 0.2 |
| | | fatty acid amide hydrolase-like | Ahyacinthus04722 | -0.8 | 0.1 |
| | | INO80 complex subunit C-like | Ahyacinthus05599 | -0.8 | 0.02 |
| | | Patched domain-containing protein 3 | Ahyacinthus17678 | -0.8 | 0.1 |
| | | glycine-rich RNA-binding protein 2-like | Ahyacinthus14948 | -0.7 | 0.1 |
| | | selenium-binding protein 1-like isoform X1 | Ahyacinthus12852 | -0.6 | 0.04 |
| | | piwi-like protein 1 | Ahyacinthus03111 | -0.4 | 0.03 |
| D × D + M × D | UP | calumenin-B-like isoform X1 | Ahyacinthus19471 | 1.6 | 0.1 |
| | | Fibroblast growth factor 1 | Ahyacinthus13035 | 1.3 | 0.2 |
| | | matrix metalloproteinase-2-like | Ahyacinthus20888 | 1.2 | 0.1 |

| | | | | |
|---|---|---|---|---|
| | glutamate dehydrogenase, mitochondrial-like | Ahyacinthus12318 | 1.0 | 0.1 |
| | predicted protein | Ahyacinthus19472 | 1.0 | 0.1 |
| | 10 kDa heat shock protein, mitochondrial-like | Ahyacinthus06892 | 0.9 | 0.1 |
| | balbiani ring protein 3-like | Ahyacinthus03095 | 0.9 | 0.02 |
| | heat shock protein HSP 90-beta-like | Ahyacinthus20571 | 0.8 | 0.1 |
| | cytochrome P450 3A8-like | Ahyacinthus12764 | 0.8 | 0.1 |
| | cytochrome b-c1 complex subunit 2, mitochondrial-like | Ahyacinthus18302 | 0.7 | 0.02 |
| | calreticulin-like isoform X1 | Ahyacinthus17491 | 0.7 | 0.1 |
| | endoplasmin-like isoform X1 | Ahyacinthus08223 | 0.7 | 0.2 |
| | endothelial differentiation-related factor 1-like | Ahyacinthus10356 | 0.7 | 0.1 |
| | hypothetical protein pdam_00000357, partial* | Ahyacinthus02976 | 0.7 | 0.01 |
| | nucleolar pre-ribosomal-associated protein 1-like | Ahyacinthus05070 | 0.7 | 0.03 |
| | FUN14 domain-containing protein 1-like | Ahyacinthus00645 | 0.7 | 0.1 |
| | coiled-coil domain-containing protein 39-like | Ahyacinthus01592 | 0.7 | 0.2 |
| | prolyl 4-hydroxylase subunit alpha-1-like isoform X1 | Ahyacinthus22060 | 0.6 | 0.1 |
| | cytochrome c oxidase subunit 6A, mitochondrial-like | Ahyacinthus01808 | 0.6 | 0.1 |
| | transmembrane protein 14C-like | Ahyacinthus02899 | 0.6 | 0.1 |
| | pyruvate carboxylase, mitochondrial-like | Ahyacinthus20647 | 0.6 | 0.1 |
| | Golgi integral membrane protein 4-like isoform X1 | Ahyacinthus03897 | 0.6 | 0.1 |
| | protein disulfide-isomerase A4-like | Ahyacinthus05582 | 0.5 | 0.02 |
| | selenoprotein K-like | Ahyacinthus03559 | 0.5 | 0.04 |
| | formin-binding protein 4-like | Ahyacinthus19849 | 0.5 | 0.03 |
| DOWN | TNF receptor-associated factor 5-like | Ahyacinthus05600 | -1.7 | 0.4 |
| | D-arabinono-1,4-lactone oxidase | Ahyacinthus08782 | -1.3 | 0.2 |
| | uncharacterized protein LOC114955431* | Ahyacinthus15775 | -1.3 | 0.1 |
| | basement membrane-specific heparan sulfate proteoglycan core protein-like | Ahyacinthus02990 | -1.3 | 0.1 |
| | TNF receptor-associated factor 5-like | Ahyacinthus05601 | -1.2 | 0.2 |
| | predicted protein* | Ahyacinthus02626 | -1.1 | 0.1 |
| | angiotensin-converting enzyme-like | Ahyacinthus01311 | -1.1 | 0.2 |
| | endothelin-converting enzyme homolog isoform X1 | Ahyacinthus04021 | -1.1 | 0.1 |
| | T-box transcription factor TBX10-like isoform X1 | Ahyacinthus02156 | -1.0 | 0.02 |
| | receptor-type tyrosine-protein phosphatase S-like | Ahyacinthus17113 | -0.9 | 0.1 |
| | Patched domain-containing protein 3 | Ahyacinthus17678 | -0.8 | 0.1 |

| | | | | | |
|---|---|---|---|---|---|
| | | ehand-like protein | Ahyacinthus13360 | -0.7 | 0.03 |
| | | actin, cytoplasmic | Ahyacinthus18280 | -0.7 | 0.04 |
| | | myotubularin-related protein 2-like | Ahyacinthus19405 | -0.6 | 0.1 |
| | | integrator complex subunit 6-like | Ahyacinthus01480 | -0.5 | 0.1 |
| | | leucine-rich repeats and immunoglobulin-like domains protein 3 | Ahyacinthus07082 | -0.5 | 0.01 |
| M × D only | UP | predicted protein* | Ahyacinthus18945 | 1.8 | 0.2 |
| | | ras-related protein Rab-30-like | Ahyacinthus13600 | 1.6 | 0.1 |
| | | cyclin-dependent kinase 2-like | Ahyacinthus06744 | 1.5 | 0.1 |
| | | KDEL motif-containing protein 1-like | Ahyacinthus15615 | 1.4 | 0.4 |
| | | uncharacterized protein LOC114955663* | Ahyacinthus13537 | 1.3 | 0.03 |
| | | protein crumbs homolog 1-like | Ahyacinthus11046 | 1.3 | 0.1 |
| | | doublesex- and mab-3-related transcription factor A2-like isoform X1 | Ahyacinthus24486 | 1.1 | 0.1 |
| | | cyclic AMP-responsive element-binding protein 3-like protein 3 | Ahyacinthus09058 | 1.0 | 0.1 |
| | | serine/threonine-protein kinase TBK1-like isoform X1 | Ahyacinthus08310 | 0.8 | 0.1 |
| | | exportin-7-like | Ahyacinthus12175 | 0.7 | 0.04 |
| | | transcriptional regulator Myc-2-like isoform X2 | Ahyacinthus03046 | 0.7 | 0.1 |
| | | ATP synthase subunit epsilon, mitochondrial-like | Ahyacinthus16123 | 0.6 | 0.04 |
| | | zinc finger protein 143-like | Ahyacinthus19867 | 0.6 | 0.01 |
| | | myosin-2 essential light chain-like | Ahyacinthus06601 | 0.6 | 0.01 |
| | | transcription initiation factor TFIID subunit 13-like | Ahyacinthus04510 | 0.6 | 0.02 |
| | | protein amalgam-like isoform X2 | Ahyacinthus24511 | 0.6 | 0.01 |
| | | unconventional myosin-VIIa-like | Ahyacinthus00047 | 0.6 | 0.02 |
| | | ubiquitin-conjugating enzyme E2 variant 2-like | Ahyacinthus16215 | 0.5 | 0.02 |
| | DOWN | uncharacterized protein LOC114977286* | Ahyacinthus17120 | -2.1 | 0.03 |
| | | uncharacterized protein LOC114961172* | Ahyacinthus10325 | -1.4 | 0.03 |
| | | histone H1-delta-like | Ahyacinthus08839 | -1.3 | 0.1 |
| | | uncharacterized protein LOC114967558* | Ahyacinthus23406 | -1.3 | 0.1 |
| | | mitochondrial glycine transporter-like | Ahyacinthus10152 | -1.1 | 0.02 |
| | | serine/threonine-protein kinase pim-1 isoform X1 | Ahyacinthus07851 | -1.0 | 0.03 |
| | | elongation factor 1-beta-like | Ahyacinthus06786 | -0.8 | 0.05 |
| | | Muscle, skeletal receptor tyrosine protein kinase | Ahyacinthus15997 | -0.8 | 0.1 |
| | | dehydrogenase/reductase SDR family member 12-like | Ahyacinthus12024 | -0.8 | 0.01 |
| | | Cell death specification protein 2 | Ahyacinthus08686 | -0.8 | 0.04 |
| | | 40S ribosomal protein S20-like | Ahyacinthus01778 | -0.8 | 0.1 |
| | | lipase member K-like | Ahyacinthus12523 | -0.8 | 0.02 |

| | | | |
|---|---|---|---|
| polyamine oxidase 1-like | Ahyacinthus14847 | -0.7 | 0.03 |
| adenosylhomocysteinase B-like | Ahyacinthus15725 | -0.7 | 0.1 |
| protein fosB-like | Ahyacinthus20539 | -0.7 | 0.1 |
| 40S ribosomal protein S12-like | Ahyacinthus02871 | -0.7 | 0.03 |
| voltage-dependent T-type calcium channel subunit alpha-1H-like | Ahyacinthus18865 | -0.7 | 0.01 |
| prostamide/prostaglandin F synthase-like | Ahyacinthus03966 | -0.7 | 0.03 |
| 40S ribosomal protein S15 | Ahyacinthus09393 | -0.6 | 0.04 |
| elongation factor 1-gamma-A-like | Ahyacinthus21246 | -0.6 | 0.02 |
| 40S ribosomal protein S9 | Ahyacinthus18437 | -0.5 | 0.05 |
| serine/threonine kinase-like domain-containing protein STKLD1 | Ahyacinthus10288 | -0.5 | 0.02 |
| translocator protein-like | Ahyacinthus12331 | -0.4 | 0.01 |

^$Log_2$ fold-change (log2FC) and standard deviation (SD) are average of *edgeR*, *limma voom*, and *DESeq2* results; *see Supplementary Table S7 for UniProt *BLASTP* results.

**Supplementary Table S7.** UniProt *BLASTP* results for the differentially expressed genes (DEGs) identified by *edgeR*, *limma voom*, and *DeSeq2* (i.e., consensus) as common and unique to *Acropora tersa* recruits from within- and between-reef crosses under heat stress relative to ambient conditions (i.e., common and unique consensus heat stress DEGs).

| Reference Annotation | | UniProt | | | | |
|---|---|---|---|---|---|---|
| **geneID** | **Description** | **Species** | **Accession** | **Description** | **Identity (%)** | **E value** |
| Ahyacinthus07208* | PREDICTED: uncharacterized protein LOC107342553 | *Acropra cervicornis* | A0AAD9Q506 | uncharacterized protein | 78.9% | 1.1E-51 |
| Ahyacinthus02416* | uncharacterized protein LOC113685661 isoform X2 | *A.cervicornis* | A0AAD9R7I3 | uncharacterized protein | 95.6% | 7.2E-78 |
| Ahyacinthus08748* | uncharacterized protein LOC114962788 isoform X1 | *Desmophyllum pertusum* | A0A9W9YCQ2 | NB-ARC domain-containing protein | 48.2% | 0 |
| Ahyacinthus15775* | uncharacterized protein LOC114955431 | *A.cervicornis* | A0AAD9V4P3 | Fibrinogen C-terminal domain-containing protein | 91.9% | 0 |
| Ahyacinthus11523* | LOW QUALITY PROTEIN: uncharacterized protein LOC114972637 | *A.cervicornis* | A0AAD9QC42 | uncharacterized protein | 85.4% | 0 |
| Ahyacinthus03761* | uncharacterized protein LOC114969772 | *A.cervicornis* | A0AAD9V1F7 | DUF4430 domain-containing protein | 70.3% | 2.5E-76 |
| Ahyacinthus17767* | PREDICTED: uncharacterized protein LOC107347891 | *A.cervicornis* | A0AAD9QY44 | uncharacterized protein | 97.8% | 2.7E-89 |
| Ahyacinthus11122* | uncharacterized protein LOC114961322 | *A.cervicornis* | A0AAD9R3G9 | uncharacterized protein | 64.7% | 9.1E-149 |
| Ahyacinthus03757* | PREDICTED: uncharacterized protein LOC107354550 isoform X1 | *A.cervicornis* | A0AAD9V1F0 | MACPF domain-containing protein | 76.2% | 0 |
| Ahyacinthus18901* | uncharacterized protein LOC114971502 | *A.cervicornis* | A0AAD9R2D0 | uncharacterized protein | 78.2% | 2.4E-55 |
| Ahyacinthus22820* | PREDICTED: uncharacterized protein LOC107337057 | *Holothuria leucospilota* | A0A9Q0YL50 | CMP/dCMP-type deaminase domain-containing protein | 36% | 4.0E-02 |
| Ahyacinthus18780* | uncharacterized protein LOC114964676 | Pocillopora damicornis | A0A3M6TEQ0 | uncharacterized protein | 27.9% | 4.9E-10 |
| Ahyacinthus12155* | uncharacterized protein LOC114964254 isoform X1 | *A.cervicornis* | A0AAD9R4J3 | Superoxide dismutase [Cu-Zn] | 83.2% | 0 |
| Ahyacinthus02729* | NA | *A.cervicornis* | A0AAD9QBK0 | uncharacterized protein | 91.6% | 1.1E-72 |
| Ahyacinthus18995* | PREDICTED: uncharacterized protein LOC107339115 | *A.cervicornis* | A0AAD9QMU4 | F-box domain-containing protein | 95.5% | 3.3E-121 |
| Ahyacinthus08657* | PREDICTED: uncharacterized protein LOC107344097 | *A.cervicornis* | A0AAD9PZD5 | uncharacterized protein | 80.9% | 0 |
| Ahyacinthus08397* | PREDICTED: uncharacterized protein LOC107348516 | *A.cervicornis* | A0AAD9R1M8 | uncharacterized protein | 93.4% | 1.1E-141 |
| Ahyacinthus15776* | PREDICTED: uncharacterized protein LOC107340658 | *A.cervicornis* | A0AAD9QFZ4 | uncharacterized protein | 94.8% | 0 |
| Ahyacinthus12545* | uncharacterized protein LOC114961810 isoform X1 | *A.cervicornis* | A0AAD9V7A3 | Aminoglycoside phosphotransferase domain-containing protein | 96.7% | 0 |
| Ahyacinthus00463* | predicted protein | *A.cervicornis* | A0AAD9QSU9 | Glypican-5 | 95% | 0 |
| Ahyacinthus18945* | predicted protein | *A.cervicornis* | A0AAD9VF13 | uncharacterized protein | 91.1% | 0 |
| Ahyacinthus13537* | uncharacterized protein LOC114955663 | *A.cervicornis* | A0AAD9UV68 | PNPLA domain-containing protein | 94.1% | 0 |
| Ahyacinthus19472* | predicted protein | *A.cervicornis* | A0AAD9V8P5 | calumenin-B | 87.4% | 0 |
| Ahyacinthus02976* | hypothetical protein pdam_00000357, partial | *A.cervicornis* | A0AAD9V5V4 | Cytochrome c oxidase subunit 6C | 97.1% | 3.7E-50 |
| Ahyacinthus10325* | uncharacterized protein LOC114961172 | *A.cervicornis* | A0AAD9QT17 | uncharacterized protein | 97.3% | 1.5E-115 |
| Ahyacinthus23406* | uncharacterized protein LOC114967558 | *A.cervicornis* | A0AAD9V550 | uncharacterized protein | 98.8% | 2.3E-116 |
| Ahyacinthus02626* | predicted protein | *A.cervicornis* | A0AAD9QBE1 | uncharacterized protein | 90.4% | 0 |
| Ahyacinthus17120* | uncharacterized protein LOC114977286 | *A.cervicornis* | A0AAD9PUC0 | GTPase IMAP family member 4 | 72.9% | 0 |

*see individual UniProt *BLASTP* result files with same geneID name.

**Supplementary Table S8.** UniProt *BLASTP* results for the seven differentially expressed genes (DEGs) identified by *edgeR*, *limma voom*, and *DeSeq2* (i.e., consensus) in all three recruit offspring groups (i.e. genotype-independent consensus heat stress DEGs).

| **Reference Annotation** | | **UniProt** | | | | |
|---|---|---|---|---|---|---|
| **geneID** | **Description** | **Species** | **Accession** | **Description** | **Identity (%)** | **E value** |
| Ahyacinthus19471* | calumenin-B-like isoform X1 | *Acropora cervicornis* | A0AAD9V914 | calumenin-B | 98.20% | 0 |
| Ahyacinthus20571* | heat shock protein HSP 90-beta-like | *A. cervicornis* | A0AAD9QU63 | HSP 90-alpha 1 | 94.50% | 0 |
| Ahyacinthus17491* | calreticulin-like isoform X1 | *A. cervicornis* | A0AAD9UYV3 | calreticulin | 89.50% | 0 |
| Ahyacinthus05582* | protein disulfide-isomerase A4-like | *A. cervicornis* | A0AAD9QZS1 | protein disulfide isomerase | 97.20% | 0 |
| Ahyacinthus19849* | formin-binding protein 4-like | *A. cervicornis* | A0AAD9QWZ5 | formin-binding protein 4 | 76.70% | 0 |
| Ahyacinthus08223* | endoplasmin-like isoform X1 | *A. cervicornis* | A0AAD9VE97 | endoplasmin/ GRP94 | 96.80% | 0 |
| Ahyacinthus17678* | Patched domain-containing protein 3 | *A. cervicornis* | A0AAD9VD06 | patched domain-containing protein 3 | 87.40% | 0 |

* see individual UniProt *BLASTP* results file with same geneID name.
